# Widespread Epistasis between Cancer Driver Mutations and Allele-Specific Copy Number Variations

**DOI:** 10.64898/2025.12.11.693698

**Authors:** Serge Merzliakov, Guanlan Dong, Andrea Castro, Mahad Bihie, Yve Nichols-Evans, Hannah Carter, Kuan-lin Huang

## Abstract

Cancer driver mutations alone are often insufficient to fully explain tumorigenesis. We demonstrate that these mutations cooperate with somatic copy number variations (CNVs) in a tissue-specific pattern of genomic epistasis. Analyzing 93,462 tumors, we identified 54 gene-cancer type pairs with significant co-occurrence of somatic mutations and CNVs. Our new Binoculars algorithm, which resolved phased DNA/RNA reads, revealed frequent preferential amplification in oncogenic mutation alleles, including AKT1 p.E17K, BRAF p.V600E, KRAS p.G12C/D/V, NRAS p.Q61K, and a fraction of gain-of-function TP53 p.R175H. Conversely, deletions selectively targeted the reference alleles, leading to loss of heterozygosity of IDH1 p.R132H and tumor suppressor mutations, including *CDKN2A* and *TP53* truncations. Lung cancer patients carrying co-occurrences of somatic mutation-CNVs in *TP53* and *KRAS* showed poorer survival than those carrying the same gene mutations. These findings reveal epistasis of cancer mutations and CNVs at an allelic resolution, suggesting specific genomic events to enhance patient stratification and therapeutic targeting.

## INTRODUCTION

The cancer genome is often characterized by a complex landscape of somatic alterations, including both somatic mutations and copy number variations (CNVs; in the purpose of this manuscript, refers to somatic CNVs), as potential drivers of tumorigenesis. To date, much of the cancer genomics research has focused on identifying driver mutations as the primary oncogenic events^1^. Numerous studies have established a catalog of recurrent mutations across various cancer types, leading to the identification of many canonical oncogenes and tumor suppressor genes. However, recent sequencing studies of non-cancerous tissues have revealed the presence of classic “driver” mutations in normal cells^2–4^, challenging the notion that mutations alone are sufficient for tumor development. These emerging findings suggest additional factors may be required in malignant progression. Recent discoveries have revealed complex interactions among genomic alterations, particularly composite mutations within the same gene^5,6^ that could synergistically enhance oncogenic potential. These findings highlight the importance of considering the epistasis between genomic alterations in disease pathogenesis.

CNVs have emerged as significant contributors to oncogenesis, particularly in malignancies characterized by low mutational burden^7^. These larger genomic alterations can lead to the amplification of oncogenes or the deletion of tumor suppressor genes^8^. Previous studies have explored the CNV-induced gene dosage effect associated with specific driver mutations in specific cancer or genes. For example, in genetically engineered KRAS G12D/+, mouse models, *KRAS* mutant alleles are selectively amplified as tumors progress^9–11^. Additional oncogenes including *KIT* and *EGFR* were also implicated by genomic studies where copy number amplifications (CNAs) preferably increase the dosage of the mutant allele^12–16^. On the other hand, interactions between somatic mutations on one allele and copy number deletions (CNDs) affecting the other allele of the same tumor suppressor gene (TSG) represent a classic example of a “two-hit” mechanism. This phenomenon, statistically formulated for *RB1* by Knudson^17^, has since been demonstrated in other cancer predisposition genes, including *ATM* and *BRCA1*^18–20^. Importantly, this two-hit mechanism is not restricted to germline variants; it also applies to somatic mutations in TSGs, where loss of heterozygosity (LOH) may occur even without copy number changes for genes including *TP53* and APC^13,21–23^. In addition to these previous reports that focus on single genes or cancer types, recent studies have expanded analyses of CNV-associated mutations to pan-cancer cohorts^7,12,13^. However, most of the documented evidence are limited at a gene level; it remains unclear which specific mutations are most affected by CNAs or CNDs in an allele-specific manner.

This study posits that the interplay between CNVs and driver mutations may not be merely additive but synergistic, mirroring the epistatic interactions observed across composite mutations. We systematically investigated the co-occurrence of somatic mutations and CNVs in cancer and evaluated their potential selection advantage. Using large-scale cancer genomics datasets from The Cancer Genome Atlas (TCGA) and AACR GENIE, comprising over 93,000 cases across 32 cancer types, we developed a robust permutation-based selection analysis that accounts for both background mutation and CNV rates to identify gene-cancer type pairs where these alterations co-occur more frequently than expected by chance. Furthermore, we incorporated phase information and analyzed the allelic imbalance of mutations influenced by CNVs at both DNA and RNA levels, assessing their impact on mutation variant allele fraction (VAF) and transcription. By leveraging this approach, we systematically identified genes and mutations with significant co-occurrence of CNVs, demonstrating tissue-specific patterns of positive selection that contributed to downstream functional and clinical consequences.

## RESULTS

### Positively Selected Co-occurrence of Mutations and CNVs in Cancer

To ascertain potential interactions between mutations and CNVs, we leveraged cancer genomics data from both TCGA^24^ and AACR GENIE^25^ cohorts, totaling 93,462 cases across 32 cancer types with availability of both genomic data types (**Figure 1A-B, Table S1A)**. We devised a permutation-based selection analysis that accounted for both background mutation and CNV rates of each tumor to determine if such co-occurrence is more prevalent than predicted solely by random chance, i.e., positively selected (**Methods**). For each cancer type, data from TCGA and each GENIE centers were separately analyzed and results combined using Edgington’s method. Our analysis revealed 54 gene-cancer type pairs with significant co-occurrence of mutations and CNVs (FDR < 0.01, **Figure 1C-D, Supplementary Figure 1A**, **Tables S1B)**, and findings across TCGA and GENIE showed agreement (r=0.37, 95% CI=0.32-0.41) for mutually evaluated pairs (**Figure 1C-D,Table S1B**).

**Figure 1.**
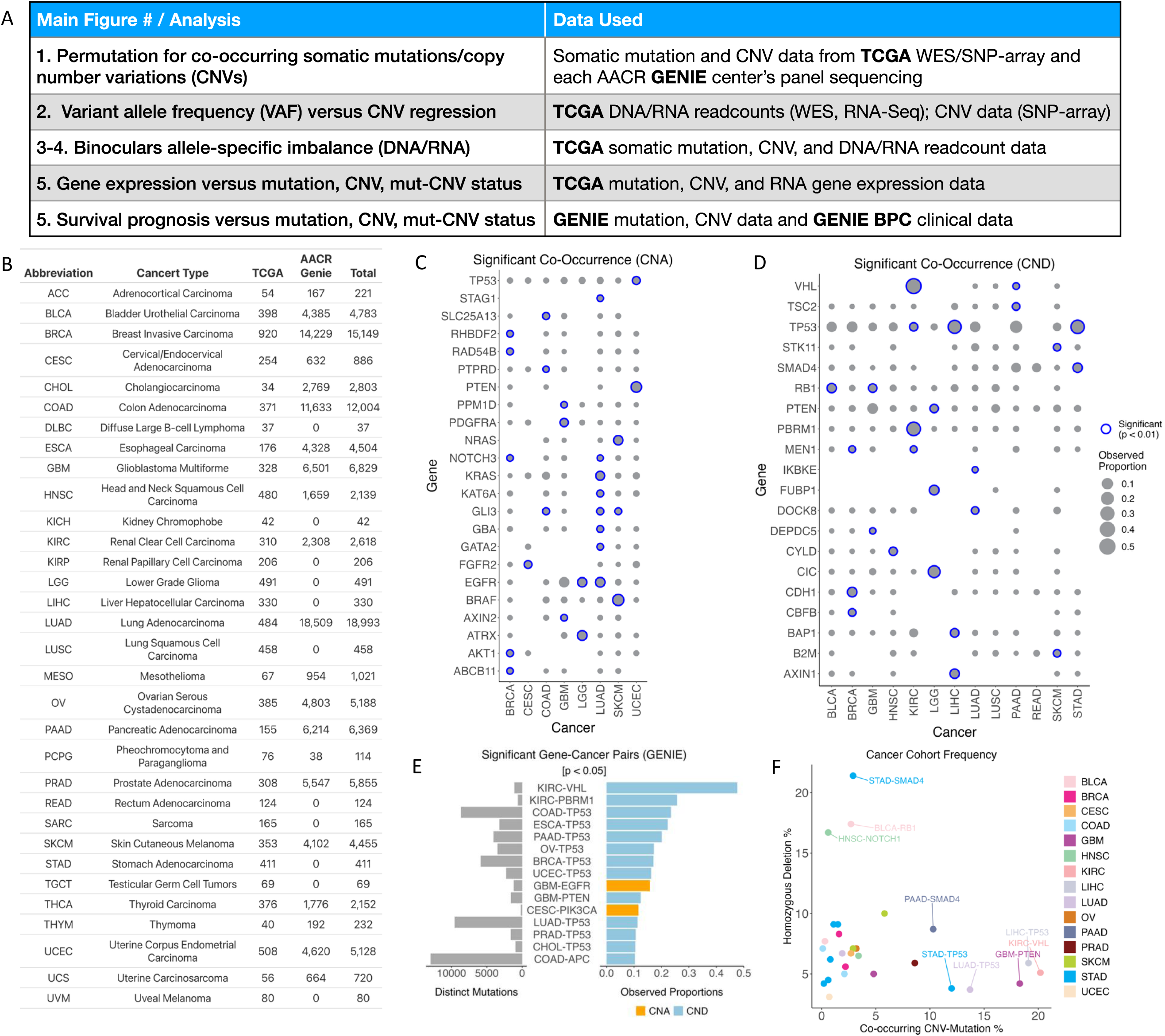
Significant Co-Occurrence of Somatic Mutations and CNVs across Cancer Types. (A) Data source for each of the main figures and analyses. (B) Cancer types and their abbreviations, with the number of cases from TCGA and AACR GENIE with both mutation and CNV data. (C-D) Significant co-occurrences of mutations and copy number deletions, including somatic (B) Copy Number Amplification (CNA) and (C) Copy Number Deletions (CND), across various cancer types, as determined by permutation analysis. Gene-cancer pair FDR values from TCGA and GENIE were summed using Edginton’s method. Observed proportions for Gene-cancer pairs were averaged across GENIE centers when multiple centers had the data on the same pair. (E) Significant gene-cancer pairs from the GENIE dataset, showing distinct mutations across all samples and observed proportion for mutation and CNV co-occurrence within the cancer cohort, where only the gene-cancer pairs with the highest proportions are shown. (F) For genes with significant mutations co-occurring with CND, we compared that frequency with the same genes’ rate of homozygous deletions—another possible mechanism of two-hits—within each of the cancer cohorts in TCGA.

Among the gene mutations that significantly co-occurred with copy number amplifications (CNAs), two genes showed association in multiple cancer types: *NOTCH3* in BRCA and LUAD, and *EGFR* in LGG and LUAD (**Figure 1C, Table S1C**). LUAD exhibited a particularly high number of 7 mutated genes with significant co-occurrence with CNAs compared to other cancer types – including *EGFR*, *GATA2*, *GBA*, *GLI3*, *KAT6A, KRAS* and *NOTCH3*. Significant co-occurrences of mutations and copy number deletions (CNDs) identified *TP53* across KIRC, LIHC and STAD (**Figure 1D, Table S1B**). In LIHC, significant co-occurrence with CNDs were observed for *AXIN1* and *BAP1* mutations. Other gene mutations strongly associated with CND included *VHL* and *PBRM1* in KIRC, *PTEN* in LGG and *SMAD4* in STAD.

Across both TCGA and GENIE datasets, *VHL* and *PBRM1* in KIRC, as well as *TP53* in various cancer types, showed co-occurring mutations and CNDs that affected the highest proportions of samples within their respectively identified cancer types (**Figure 1D-E, Supplementary Figure 1A, Tables S1E**). Oncogenes affecting high proportions of samples include *BRAF* in SKCM, *EGFR* in GBM, and *PIK3CA* in CESC (**Figure 1C,E, Table S1D**). Most of the identified gene-cancer type pairs showing strong co-occurrence and high frequencies were consistent between TCGA and GENIE, demonstrating the robustness of the identified genes (**Tables S1C-E**). These results provide evidence for wide-spread positive selection of co-occurring mutations and CNVs in a gene- and tissue-specific manner.

The mechanistic origin of how mutations could be positively selected by CNVs remains unclear. Extra chromosomal circular DNA (eccDNA) has emerged as a potential cancer driver associated with high oncogene expression^26,27^. To assess whether CNA-associated mutations may originate from eccDNA, we analyzed mutations from significant gene-cancer type pairs associated with CNAs (n=13). There were 2,923 samples with mutation-CNA co-occurrence, among which 338 samples had eccDNA fragments large enough to contain a full copy of a significant gene (**Supplementary Figure 1D, Table S1E**) (Note: 360 TCGA samples had eccDNA calls based on the ATAC-seq data^28^). We found 6 significant gene-cancer type pairs impacted by CNAs where samples also contained eccDNA copies of the gene (**Supplementary Figure 1C, Table S1F**), including *PIK3CA* in BRCA, *KRAS* in COAD and LUAD, *BRAF* in SKCM, and *EGFR* in LUAD. However, the proportions of samples where this occurred were very low—the highest being 1.2% of samples of *PIK3CA* in BRCA and *KRAS* in COAD—and *PIK3CA* being the gene showing the highest counts of co-occurring CNV/mutations overlapped with eccDNA (N=4). Thus, eccDNA did not fully explain the origin of oncogenic mutations accompanied by amplifications.

While co-occurring mutation-CNVs represent one type of two-hit event, genomic two-hits could also happen when both copies of the gene were deleted (homozygous deletions^29^). We compared the frequency of these genes affected by co-occurring mutation-CNVs and homozygous deletions^30^ in each of the cancer types (**Figure 1F, Table S1G**). This analysis identified different TSG classes, including those (1) predominantly affected by homozygous deletions, including *SMAD4* in STAD, *RB1* in BLCA, and *NOTCH1* in HNSC; (2) predominantly affected by co-occurring mutation/CNDs, including *TP53* in LIHC/LUAD/STAD, *VHL* in KIRC, and *PTEN* in GBM (**Figure 1F, Table S1G**). It is possible that the different preferences of two-hit mechanisms could be correlated with essential genes or other possibly co-deleted tumor suppressor genes in the adjacent chromosomal region^31^.

### Phase and Allelic Imbalance in Co-occurring Mutations and CNVs

While previous analyses demonstrate the enrichment of co-occurring mutations and CNVs, they do not capture allele-specific effects through phase information, i.e., whether the identified CNVs specifically affect the mutant or the reference allele. By obtaining read count data from TCGA, we determined the effect of CNVs on mutation VAF of the affected gene across tumors to distinguish the four scenarios that may arise: (1) Increased mutation VAF in the CNA-affected gene implies preferred amplification of the mutant allele. (2) Increased mutation VAF in the CND-affected gene implies preferred deletion of the reference allele. (3) Decreased mutation VAF in the CNA-affected gene implies preferred amplification of the reference allele. (4) Decreased mutation VAF in the CND-affected gene implies preferred deletion of the mutant allele. To identify significant associations between CNA/CND and mutation VAF across genes in each cancer type, we used a multiple regression model with adjustments for tumor purity and clinical variables (**Methods**). A significant allelic imbalance could only be identified when sufficient statistical power is afforded by clonality of both events as well as the sequence coverage of the allele under investigation, and in the cases of CNAs, when the somatic mutation occurs *before* an amplification. When evaluating a single mutation found across samples, a mutant allele that shows consistent directions of allelic imbalance offers cross-validation for such signal.

From a total of 1,789 gene-cancer type contexts pairs with sufficient data for analysis, we discovered 15 associations where a gene’s CNAs and CNDs were significantly linked (FDR < 0.05) to alterations in mutation VAF across tumors in a given cancer type (**Figure 2A-B, Table S2A-B**). Nearly all identified CNA (with the exception of *SPTA1* in UCEC) were associated with increased mutation VAFs, which suggests the mutations in these genes were amplified by copy number changes. These included some notable oncogenes known to be drivers in the found cancer types, such as *PIK3CA* in BRCA, *BRAF* in SKCM, and *EGFR* in LGG and LUAD, and *KRAS* across several cancers including PAAD, LUAD and COAD (**Figure 2C**, **Table S2C**). We also identified an emerging driver *CTNNB1* in LIHC. For the 4 CND associations (FDR < 0.05), all linked mutations exhibited increased VAF, including *TP53* in SARC/PAAD and *ATM* in BLCA (**Figure 2D, Table S2D**). This data supports a “two-hit” scenario where one copy of the gene was disrupted by mutations and the other by deletions.

**Figure 2.**
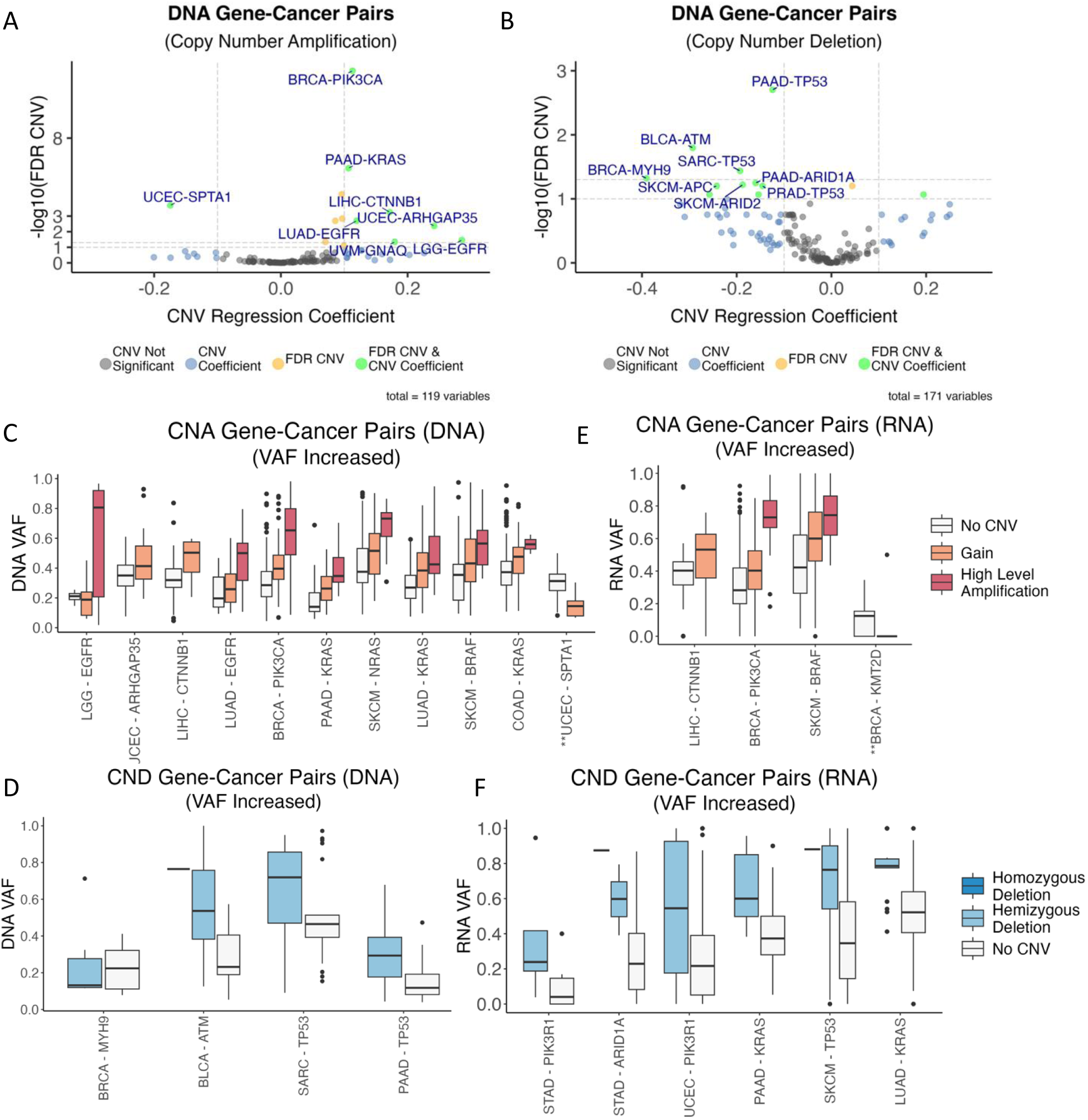
Impact of Co-occurring CNV on DNA- and RNA-level Variant Allele Frequency (VAF) (A) The effects of CNV on DNA VAF of the co-occurring mutations as shown by volcano plot based on regression coefficients and statistical significance (FDR) for Copy Number Amplification (CNA) in gene-cancer type pairs. (B) The effects of CNV on DNA VAF of the co-occurring mutations as shown by volcano plot based on regression coefficients and statistical significance (FDR) for Copy Number Deletions (CND) in gene-cancer type pairs. For (A-B), the x-axis represents CNV regression coefficients, and the y-axis represents the -log10(FDR). Gene-cancer type pairs with FDR<0.1 and CNV coefficient pass thresholds are highlighted in green, only FDR significant pairs in orange, and CNV coefficient significant pairs in blue, whereas only those with FDR<0.05 were plotted in panels C-F. FDR cutoffs at 0.1 and 0.05 levels are drawn. (C-F) Significant gene-cancer type pairs (FDR<0.05) for which CNVs were associated with alterations in DNA and RNA VAFs of the co-occurring mutations and their VAF distribution by CNV types, including copy number gain, high amplification, hemizygous deletion, and homozygous deletion, as defined by the GISTIC algorithm. The plots show the DNA VAFs impacted by (C) CNA and (D) CND and RNA VAFs impacted by (E) CNA and (F) CND. In (C) and (E), the two associations marked with ’**’ had a decreased VAF in the associated gene mutations; all other associations showed increased mutant VAFs.

In addition to DNA-level analyses, we investigated the influence of CNVs on mutation VAFs at the mRNA level across tumors. Out of 1,515 gene-cancer type pairs with sufficient data for analysis, we identified 10 significant associations (FDR < 0.05) at the RNA level (**Figures 2E-F, Table S2E-F**). Three out of four CNA associations showed an increased RNA VAF, including *CTNNB1* in LIHC, *PIK3CA* in BRCA and *BRAF* in SKCM. Only *KMT2D* in BRCA had a decreased RNA VAF (**Figures 2E, Table S2E**). For CND associations, all six showed increased RNA VAFs of the linked mutations, suggesting copy number deletion of the reference allele in these genes. We identified *PIK3R1*—a tumor suppressor down-regulating the PI3K pathway showing this two-hit scenario in both STAD and UCEC. *KRAS* deletions were associated with increased RNA VAFs of *KRAS* mutations in both PAAD and LUAD (**Figures 2F, Table S2F**), suggesting that when deletions occurred at this locus, the *KRAS* mutant allele is preferentially retained and expressed.

We next sought to directly link RNA allelic imbalance of these mutations to their respective gene’s total expression in each cancer type. In CNA-affected genes, particularly *BRAF* in SKCM and *PIK3CA* in BRCA, higher RNA VAF showed trends of positive correlation with total gene expression (**Supplementary Figures 2A–D, Table S2G**), albeit not significant after multiple testing corrections. For CND-affected genes (**Supplementary Figures 2E–H, Table S2G**), trends of potential positive correlations were seen for *TP53* in SKCM and *KEAP1* for LUAD, but there were no significant correlations.

### Allelic Imbalance of Individual Mutations that Co-occur with CNV

Having established that gene mutation VAFs were linked to CNVs across tumors, we next sought to identify individual mutations that showed significant allelic imbalance in these scenarios, which would indicate the selection of the mutant allele by tumor cells via CNVs at an intra-tumor level. We devised an algorithm, Binoculars (Binomial test for co-occurring mutation and CNV pairs), that conducts a best-matched binomial test for each of these mutations observed in each TCGA individual using a background DNA VAF that maximizes matching for each sample, gene, and/or cancer type, in order to more accurately account for sample purity and background selection rate (**Methods**). This rigorous procedure will likely result in false negatives, as power is limited by read depth and multiple testing burden obtained for the single mutation in a given sample. Nonetheless, Binoculars effectively identified specific, recurrent CNA-impacted and CND-impacted mutations that showed significant allelic imbalance across cancer types (**Figure 3A-C, Table S3A-C**).

**Figure 3.**
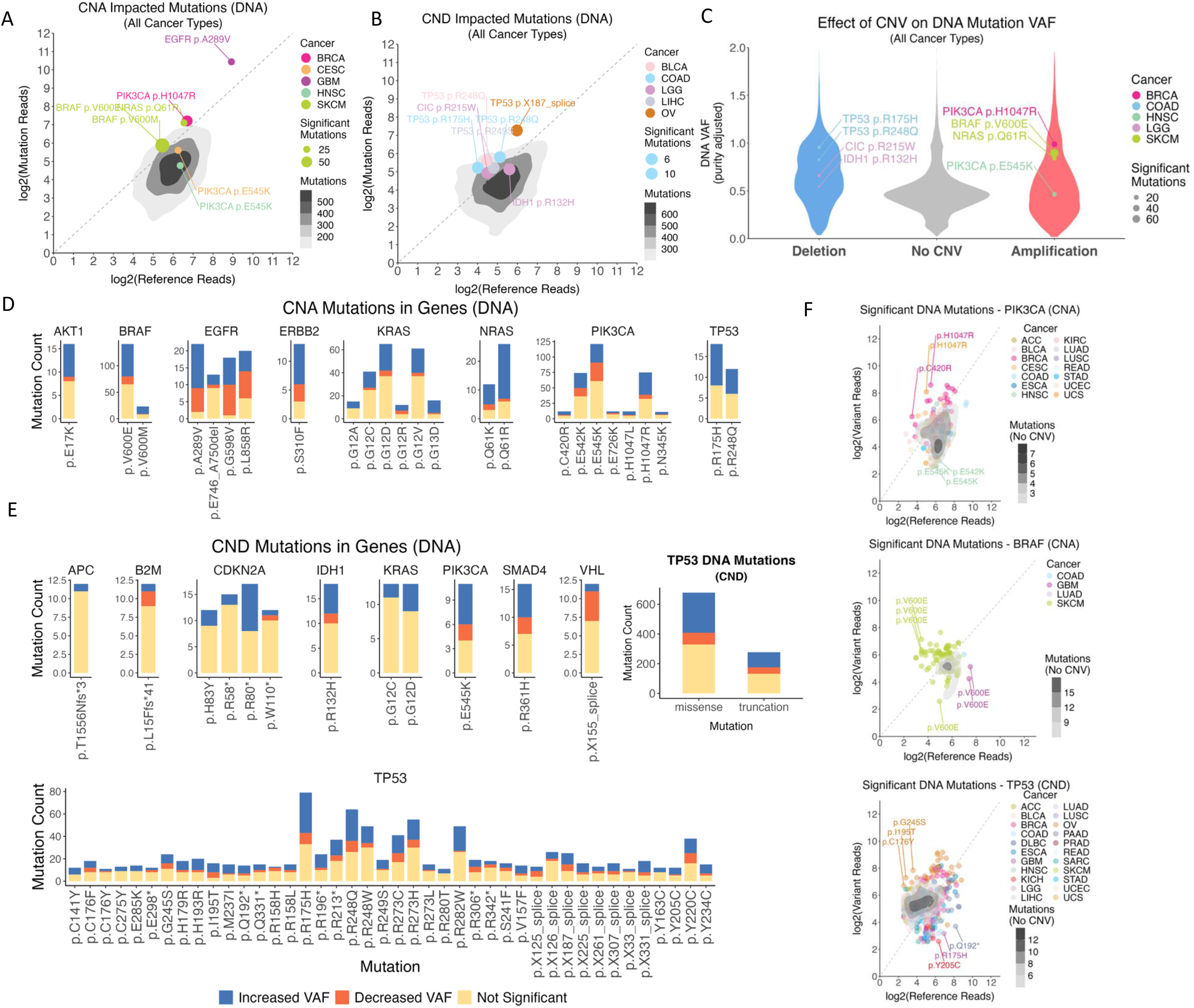
DNA-level Allelic Imbalance of Somatic Mutations Co-occurring with CNA or CND. (A-B) Density plots of mutation alleles significantly impacted by (A) CNAs and (B) CNDs across all cancer types and samples, as shown by their log2 counts of mutation and reference allele DNA read counts. The most common significant mutations are labelled and color-coded by cancer type, with the size of the point indicating the number of mutations across all samples for a single cancer type. The labelled mutations read counts (mutation and reference) are mean values for a single cancer type. The diagonal line indicates an equal amount of reference and mutant allele in a pure heterozygous state, albeit the mutant allele reads are likely diluted by tumor purity. (C) Violin plots of the mutations’ purity-adjusted DNA VAFs by their copy number status, with the most frequent mutations labelled. (D-E) The most frequent mutations with associated (C) CNA and (D) CND across all cancer types and samples with at least 10 mutations, showing proportions of mutations which significantly impacted VAF at the DNA level, including increased (Blue) or decreased (Orange) VAF and those which did not (Yellow). TP53 mutations co-occurring with copy number deletions are further grouped by mutation type. (F) Examples of gene mutations whose DNA VAFs were impacted by CNV, plotted where each dot denotes a single mutation in a single tumor, including PIK3CA, BRAF and TP53 mutations co-occurring with CNAs or CNDs. Density plots show VAF for mutations without CNVs.

Most of the CNA-impacted mutations with significant allelic imbalance exhibited increased DNA copies of the mutant allele, as heterozygous mutations would show <50% DNA VAF after accounting for tumor purity (**Figure 3A, Table S3A**). Oncogenes such as *BRAF*, *EGFR*, *KRAS*, and *PIK3CA* harbored the highest count of significant mutations. Binoculars identified specific mutations frequently showing significant allelic imbalance in many affected tumors; for instance, 75 of the significant *BRAF* p.V600E mutations, predominantly in SKCM, showed a mean tumor VAF of 0.57 suggesting strong selections of the mutant allele via amplification. We next examined the mutations with the highest counts of allelic imbalance among the genes with the most CNA-impacted mutations, counting how often each mutation’s VAF was decreased vs. increased (**Figure 3D, Table S3D**). For *AKT1*, *BRAF*, *EGFR*, *NRAS*, and *TP53*, the majority of mutations co-occurring with amplifications showed significantly increased VAF compared to the respective gene’s mutations in tumors without co-occurring CNVs (**Supplementary Figure 3**). CNA-impacted mutations in *NRAS* and *KRAS*, including p.G12A, p.G12C, p.G12D, p.G12R, p.G13D, p. Q61K, and p. Q61R, were predominantly associated with increased VAF, where their positive selection by amplifications aligns with previous studies^32^. Similarly, *PIK3CA* gene mutations p.E542K, p.E545K, and more particularly p.H1047R, showed higher fractions of the mutant allele being amplified by CNV. *EGFR* mutations at p.A289V, p.G598V, and p.L858R showed slightly high fractions of mutations with significant allelic imbalance. Interestingly, while most of the significant TP53 p.R175H mutations co-occurred with deletions that led to LOH, we also found several incidences where *TP53* amplifications are associated with increased mutant allele p.R175H, which is consistent with TP53 p.R175H being a likely gain-of-function mutation capable of modulating TP53’s interaction with other proteins^33^. The amplifications associated with increased mutation VAF likely indicate the preference and positive selection of the mutant allele during tumor development.

Most CND-impacted mutations identified as significant by Binoculars also showed allelic imbalance where the mutant allele has significantly higher counts than the reference allele (**Figure 3B, Table S3B**). The majority of these mutations affected tumor suppressor genes, aligning with the two-hit hypothesis. CND-associated *TP53* mutations were by far the most common, including TP53 p.R248Q, p.282W, p.273H, p.248W, p.220C, p.273C, and p.213*, which are likely reflecting two-hits affecting *TP53*. Other gene mutations affected by CNDs included CDK2NA p.R80*, IDH1 p.R132H, and SMAD4 p.R361H, all showing the preferential deletion of the reference allele for potential two-hit effects (**Figure 3E, Table S3E**). Interestingly, two *IDH1* monomers are required to form a catalytic homodimer; the deletion-induced LOH associated with IDH1 p.R132H will result in inactive dimers, rather than the wild-type/R132H dimers that exhibit neomorphic activity^34^. SMAD4 p.R361H is a hypomorph mutation for which a small molecule has been derived to rescue *SMAD4*-mediated transcriptional activity^35^, and its coupled deletion will likely further reduce *SMAD4* functional level. Overall, these examples highlighted the effect of intra-tumor selection for these mutations via CNV, dominated by mutant allele amplification or deletion of the wild type allele.

### Transcriptional Impact of Co-occurring Mutations and CNVs

Using Binoculars, we also examined the RNA allelic imbalance of each mutation impacted by CNA or CNDs at the mRNA level within the tumor sample, finding individual mutations showing significant allelic imbalance (**Figure 4A-C**, **Tables S4A-B**). The genes harboring the most significant mutations co-occurring with amplifications (**Figure 4D, Table S4C**) mirrored those identified in the DNA-level allelic imbalance analysis, including known oncogenes such as *PIK3CA*, *EGFR*, *BRAF*, and genes of the RAS family. Overall, there was a stronger trend for increased proportions of CNA-impacted mutations showing increased mutant RNA VAF, in comparison with DNA-level results. This RNA-level allelic imbalance was especially strong for *AKT1*, *EGFR* and *TP53* mutations, where nearly all mutations showed significantly increased RNA VAF, suggesting these co-occurring mutations may be preferentially transcribed in addition to being preferentially amplified.

**Figure 4.**
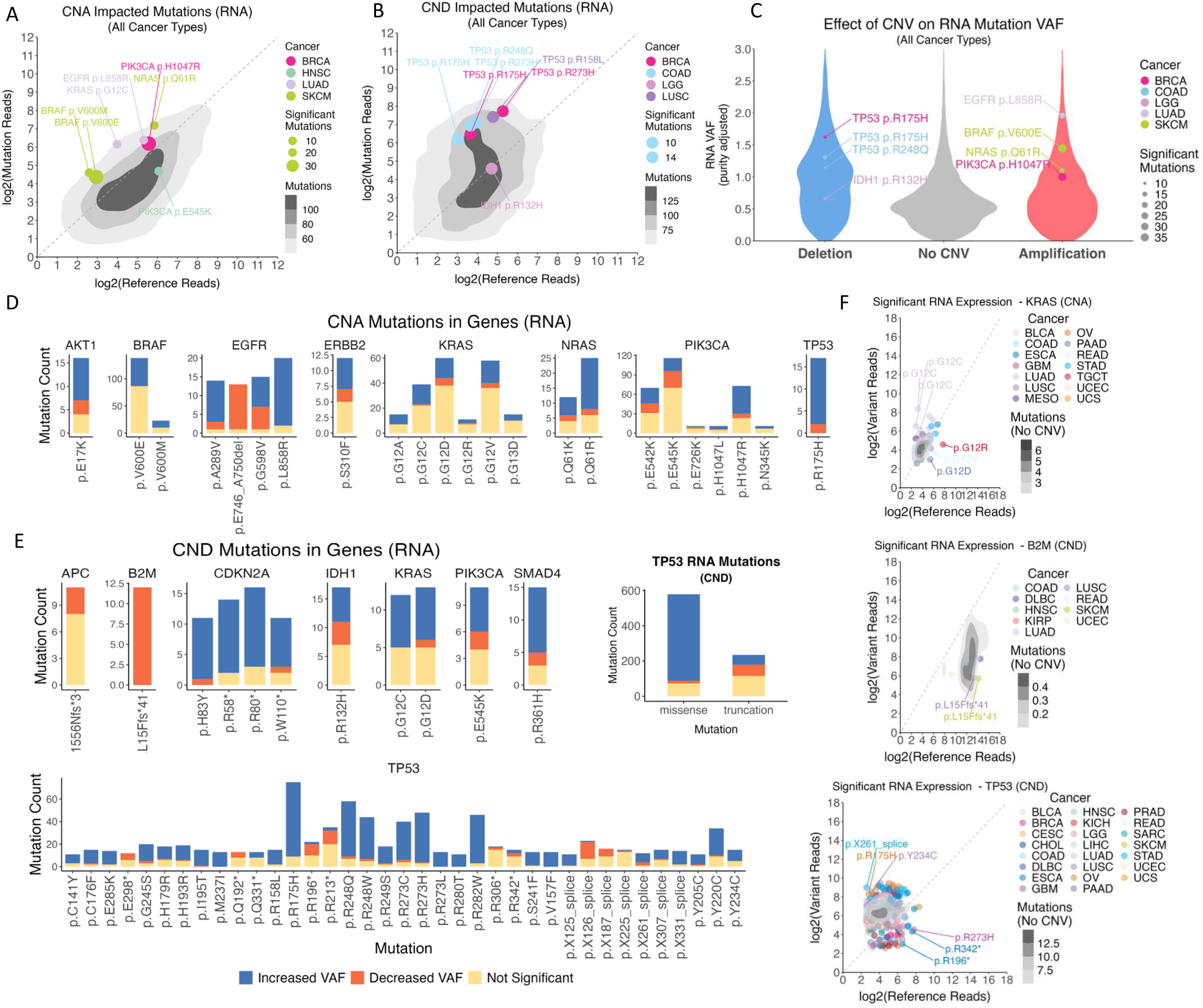
RNA-level Allelic Imbalance of Somatic Mutations Co-occurring with CNA or CND. (A-B) Density plots of mutation alleles significantly impacted by (A) CNAs and (B) CNDs across all cancer types and samples, as shown by their log2 counts of mutation and reference allele RNA read counts. The most common significant mutations are labelled and color-coded by cancer type, with the size of the point indicating the number of mutations across all samples for a single cancer type. The labelled mutations read counts (mutation and reference) are mean values for a single cancer type. The diagonal line indicates an equal amount of reference and mutant allele in a pure heterozygous state, albeit the mutant allele reads are likely diluted by tumor purity. (C) Violin plots of the mutations’ purity-adjusted RNA VAFs by their copy number status, with the most frequent mutations labelled. (D-E) The most frequent mutations with associated (C) CNA and (D) CND across all cancer types and samples with at least 10 mutations, showing proportions of mutations which significantly impacted VAF at the RNA level, including increased (Blue) or decreased (Orange) VAF and those which did not (Yellow). TP53 mutations co-occurring with copy number deletions are further grouped by mutation type. (F) Examples of gene mutations whose RNA VAFs were impacted by CNV, plotted where each dot denotes a single mutation in a single tumor, including KRAS, B2M and TP53 mutations co-occurring with CNAs or CNDs. Density plots show VAF for mutations without CNVs.

Binocular analysis of CND-associated mutations at the RNA level (**Figure 4E, Table S4D**) revealed that *TP53* mutations were overwhelmingly predominant, consistent with our findings at the DNA level. *TP53* mutations, including p.R175H, p.R248Q, p.R273H, p.R282W, p.R248W, p.R273C, and p.R273H, represented the majority of significant mutations showing RNA allelic imbalance. We also identified tumor suppressor genes whose deletions, surprisingly, lead to an allelic imbalance of higher ratio of reference allele suggesting a lack of LOH, including *B2M* and *APC* truncations (**Figure 4E, Table S4D**) (**Supplementary Figure 4**). The B2M p.L15Ffs*41 is a frameshift deletion affecting the *B2M* protein with 119 amino acid (a.a.) residues. Since *B2M* encodes for a protein vital for the structural integrity of MHC Class I complexes, any non-functional B2M subunits would disrupt the presentation of tumor antigens to Cytotoxic T cells. The APC T1556Nfs*3 affects APC protein that is 2,843 a.a. in length. The more apparent allelic imbalance for these two mutations at the RNA compared to DNA level suggests that non-sense mediated decay (NMD) is likely further responsible for the diminished mutant transcript. Our findings reveal that these genes may act as haploinsufficient drivers in cancer where an intact copy of the gene is still preferred. For example, while *B2M* loss eliminates MHC I presentation to evade T cell recognition, it simultaneously removes inhibitory ligands for NK cells, potentially triggering NK-mediated cytotoxicity that could be disadvantageous to tumors. CDKN2A truncations (p.H83Y, p.R58*, and p.R80*; but not the p.W110* closer to the canonical stop-codon) had almost all of significant mutations where the deletions’ co-occurrence was associated with increased RNA VAF of the mutant allele, suggesting the deletion of the wild-type allele and two-hit scenarios consistent across DNA- and RNA-allele analyses. Intriguingly, when CNDs occur at oncogene loci, they typically are selected for deletion of the reference allele rather than the mutated KRAS p.G12C/D or PIK3CA p.E545K allele (**Figure 3E**, **Figure 4E**).

To determine if the effects of CNVs on DNA VAF are consistent with RNA VAF, we analyzed each mutation with at least 10 samples with DNA/RNA VAF data. Many oncogene mutants co-occurring with CNAs, including BRAF p.V600E, EGFR p.A289V, KRAS p.G12V/Q16R, and PIK3CA p.E542K/E545K/H1047R, showed strong positive correlations (FDR < 0.05, Pearson Correlation) (**Supplementary Figures 5A-H**, **Table S5A-L**). Among the four tested *TP53* mutations associated with CNDs, DNA/RNA allelic imbalance analyses consistently showed increased mutation VAF suggesting LOH. However, RNA VAFs for certain mutations of p.R175H/R273C/R282W/R248Q show further RNA-level imbalance beyond their DNA VAFs, suggesting additional transcriptional regulation may affect these mutations (**Supplementary Figures 5I-L**).

As many CND-associated *TP53* mutations showed DNA and RNA level imbalances, we also stratified *TP53* alterations impacted by CND by missense versus truncating mutations (**Figures 3E, 4E**). At the DNA level, 40% of missense mutations and 37% of truncation mutations showed increased VAF, and much less showed decreased VAF. In comparison, at the RNA level, 85% of missense mutations and 23% of truncation mutations showed increased RNA VAF. This suggests that in addition to the preferential genomic deletion of the *TP53* reference allele, additional transcriptional regulatory mechanisms may further upregulate the RNA expression of the *TP53* missense mutations when the wild type allele is deleted.

To identify genes where co-occurring mutation and CNV show synergistic effects on gene expression, we took a two-step approach: (1) We applied AeQTL^36^ to identify, for each cancer type, genes whose mutations are associated with their expression after adjusting for covariates. (2) For significant gene-cancer type pairs, a linear regression model was fitted to include mutation status, CNV (considered as a continuous GISTIC^37^ value between -2 and 2), interaction between mutation status and CNV, and clinical covariates. For missense mutations, we identified 15 gene-cancer type pairs showing significant interaction effects between CNV and mutation status (FDR<0.05)(**Supplementary Figure 6A-B, Table S6A-B**). Positive interaction effects, where the co-occurrence of mutations and CNVs were associated with higher gene expression than either mutation or CNV alone, were observed in multiple known proto-oncogenes, including *PIK3CA* in LUAD, *KIT/HRAS/BRAF* in SKCM, *JAK3* in LUSC, *AKT1* in PRAD, and *EGFR* in COAD and LGG. While less observed for missense mutations, we also identified negative interaction effects, where the co-occurrence of mutations and CNVs were associated with lower gene expression than either mutation or CNV alone, for *HRAS* in BRCA, *CTNNB1* in CESE, *TP53* in READ, and *CDKN2A* in SKCM.

On the other hand, while most truncating mutations alone showed a negative effect on gene expression as expected from NMD, interaction effects between CNV and mutation status varied across genes (**Supplementary Figure 6A, Table S6B**). *TP53* and *NF1* showed negative interaction effects across most cancer types. Of 21 gene-cancer type pairs with significant interaction effects (FDR<0.05), 14 showed positive interaction effects with the largest effects found in *FGFR3* in BLCA, *CDK12* in STAD, and *MET* in LGG (**Supplementary Figure 6B, Table S6B**). Co-occurring truncating mutation and CNV can also exhibit opposite interaction effects in different cancers. For instance, the interaction effect in *BCOR* is highly negative in ESCA but positive in COAD. In some cases, interaction between mutation status and CNV is consistent between missense and truncating mutations for certain genes, such as *EGFR* in LUAD which showed a positive interaction effect for both. However, *TP53* mutation and CNVs in LUSC has the opposite interaction effects which are positive for missense mutations but negative for truncating mutations (**Supplementary Figure 6C, Table S6B**), suggesting that the synergistic impacts between mutation and CNV may be mutation type and context specific.

### Clinical Prognosis of Patients Carrying Co-occurring Mutation and CNVs

Lastly, we investigated the effects of mutation and CNV co-occurrence on patient survival using curated clinical data from the AACR Project GENIE Biopharma Collaborative (BPC), for which the data is available for lung and colorectal cancers, recognizing that prognostic associations found in such cohort require validation in prospective and clinical trial cohorts with better matched case and control samples. In non-small cell lung cancer, mutations and CNVs in *EGFR*, *KRAS* and *TP53* were the most common, so we focused on the prognosis of patients stratified by carrying these respective gene’s mutations only, CNV only, and co-occurring CNV and mutations using a Cox proportional hazards model that included key clinical covariates (**Methods**). In patients with co-occurring *TP53* mutation and CNV, survival times were reduced compared to those showing *TP53* mutation alone (**Figure 5A**, **Table S6C**, p=0.003, HR 1.5, CI 1.15-1.96). Comparable results were found for *EGFR* (**Figure 5B, Table S6D,** p=0.006, HR 1.56, CI 1.14-2.15) and *KRAS* (**Figure 5C**, **Table S6E**, p=0.02, HR 1.5, CI 1.07-2.12). For patients with only CNVs for *EGFR* but not *KRAS*, survival times were also significantly reduced compared to patients carrying only mutations (**Figure 5B, Tables S6D**). We also compared the survival of lung cancer patients with co-occurrence of *TP53*, *KRAS* and *EGFR* mutation and CNVs to that of all other patients without each of these mutations/CNVs, respectively, and found similar results (**Supplementary Figure 7A-C, Table S7A-C**).

**Figure 5.**
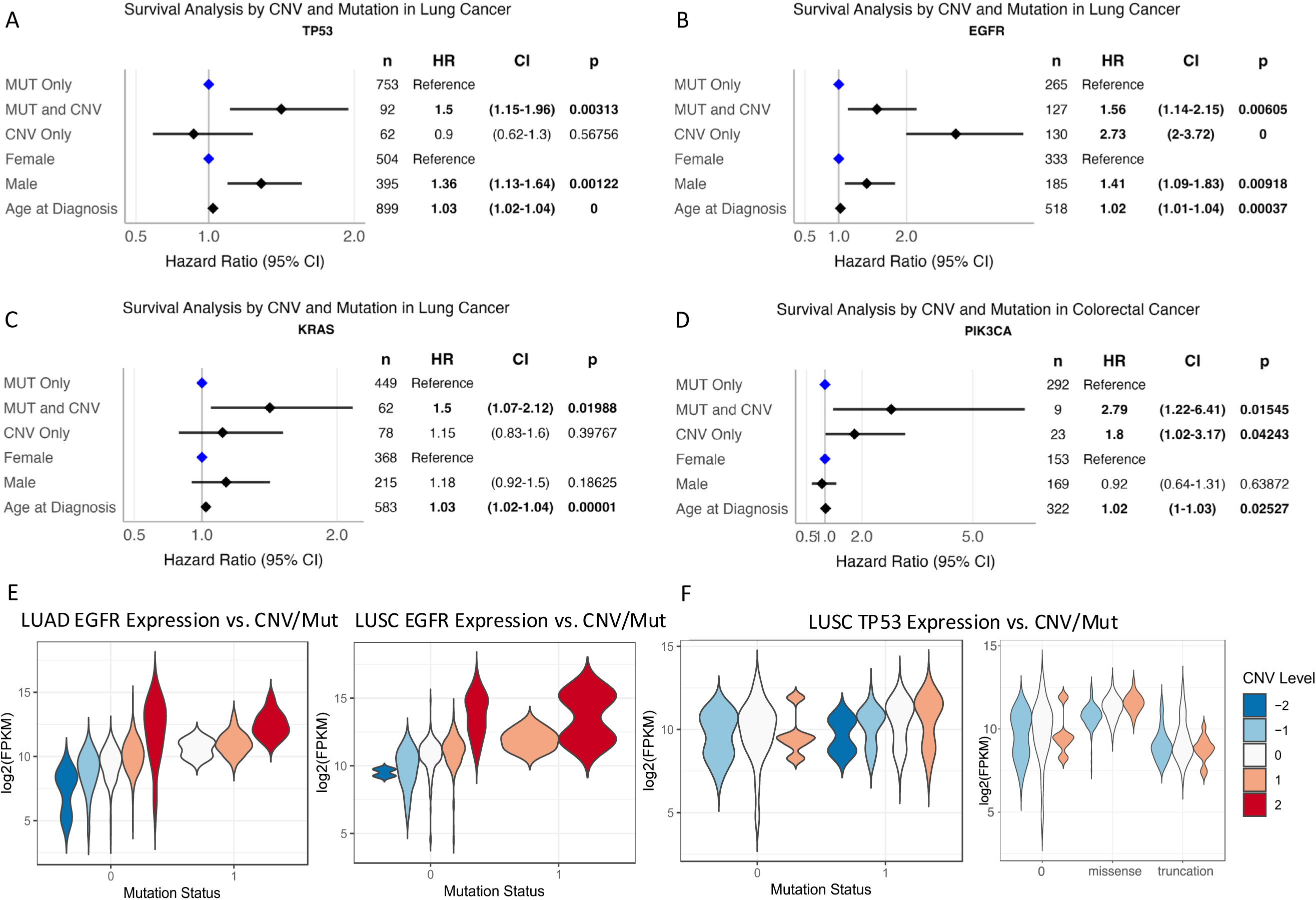
Clinical Prognosis of Carriers Stratified by Gene Mutation, CNV, and Co-occurring CNV/mutations in Lung and Colorectal Cancer. (A)-(D) The survival prognosis of stratifying patients by mutation only, CNV, and co-occurrence of CNV/mutations, as shown by forests plots in multivariate Cox regression models corrected for key demographic variable, as shown for (A) *TP53*, (B) *EGFR*, (C) *KRAS* for lung cancer and (D) *PIK3CA* for colorectal cancer in the AACR Genie BPC dataset. The reference hazard level is set to one for patients carrying mutations only in the specified gene. **Cancer Stage was a covariate in the model but is not shown in the figures. (E) Differential expression of *EGFR* by mutation and CNV status in the TCGA LUAD and LUSC cohort. (F) Differential expression of *TP53* by mutation and CNV status in the TCGA LUSC cohort.

We reasoned this may be due to dosage effects, where patients carrying only mutations vs. those with co-occurring mutations/CNVs show different levels of gene expressions. Indeed, *EGFR* showed positive interaction effects for mutations and CNVs in multiple cancer types (**Supplementary Figure 6**), including LUAD (**Figure 5E, Table S6A**). Interestingly for *TP53*, the differences between CNV-mutation subsets versus those carrying only CNVs or mutations were not found when considering all *TP53* mutations, where *TP53* mutation carriers show a bimodal pattern of expression. However, when stratifying *TP53* carriers by truncations (which likely lead to two-hit) vs. missense (which may result in gain-of-function), missense carriers show a synergistically higher *TP53* gene expression levels whereas truncation carriers showed a lower *TP53* expression compared to those without *TP53* mutations (**Figure 5F, Table S6B**). These results demonstrated that stratifying patients by mutation-CNV co-occurrence may provide additional prognostic value for patients carrying *TP53* and *KRAS*, whereas patients carrying *EGFR* CNVs may already show worse prognosis similar to those carrying co-occurring mutation-CNV of these genes.

We also evaluated the prognosis of lung cancer patients treated with *EGFR* tyrosine kinase inhibitors (TKI) and monoclonal antibodies (MAB) and whether in this subgroup, co-occurrence of CNVs and mutations affected prognosis compared to those having *EGFR* mutation alone. The survival analysis was stratified by age as survival impacts of monoclonal antibody treatments differed for younger (age < 60) vs. older patients. However, the current AACR BPC study may still yet provide sufficient sample size to reach robust conclusion in this stratified analysis (**Supplementary Figure 7D-E, Table S7D-E)**.

For colorectal cancer, we were able to analyze the effect of mutation-CNV co-occurrence on patient prognosis for three genes, namely *APC*, *FBXW7*, and *PIK3CA*. Only *PIK3CA* showed significant association, where there was strong evidence of poorer survival in patients with mutation-CNV co-occurrence (p=0.035, HR 2.38 with CI of 1.06-5.33) (**Figure 5D, Table S7G**). Similarly, comparing the survival of colorectal cancer patients with *PIK3CA*, *FBXW7* and *APC* mutations/CNVs to all other patients without each of these mutations, only patients with concurrent *PIK3CA* mutations and CNVs had poorer survival outcomes (**Supplementary Figure 8, S8D-E**).

The current precision medicine paradigm considers stratifying patients by mutation status in driver genes. Here we show that stratification of patients carrying mutations, CNVs, and co-occurring mutations and CNVs may identify patient subsets showing different prognoses due to mechanisms that may be associated with the aggressiveness of the tumor, e.g., two-hits affecting *TP53* or amplification of the mutant *PIK3CA* alleles, that may be worthy of consideration in clinical stratification.

## DISCUSSION

Our study provides a systematic catalog for the co-occurrence of mutations and CNVs in a mutation-and cancer-type-specific manner. Our results show that certain mutations, when accompanied by CNVs, exhibit non-random, positively selected interactions in tumorigenesis. By leveraging genomic data of over 93,000 tumors from TCGA and AACR GENIE, we identified 54 associations where a gene’s mutations and CNVs co-occur more frequently than expected by chance. This suggests a positive selection for their co-occurrence, providing a potential explanation for why non-cancerous tissues often harbor classic “driver” mutations in normal cells^2–4^ as additional selections by CNVs may be needed.

The co-occurrence of CNAs with oncogene mutations, such as *EGFR*, *PIK3CA*, *KRAS*, and *NRAS*, highlights their role in enhancing tumorigenic potential by amplifying mutant alleles. In contrast, CNDs associated with tumor suppressor genes, such as *TP53*, *CDKN2A*, and *SMAD4*, support the classic two-hit hypothesis^17^ where a loss-of-function (LOF) mutation of one allele and deletion of the other via LOH contributes to tumorigenesis, but do not apply to certain *B2M* and *APC* truncations. While previous studies have identified the co-occurrence of mutations and CNVs affect certain oncogenes and tumor suppressors, they do not focus on pinpointing specific mutations of interest^8,21^. By considering the phase of CNVs in relation to mutations, we showed that mutations co-occurring with CNAs generally result in increased VAF, signifying preferential amplification of the mutant allele. Similarly, CNDs often lead to increased VAF of tumor suppressor mutations, consistent with the deletion of the wild-type allele. Our newly developed Binoculars algorithm enabled precise analysis of phased DNA and RNA read counts for a given tumor’s mutation. This approach uncovered preferential amplification of key mutant alleles, such as AKT1 p.E17K, BRAF p.V600E, KRAS p.G12C/D/V, NRAS p.Q61K, and the gain-of-function TP53 p.R175H. Conversely, reference alleles in genes affected by loss-of-function mutations, such as truncations in *CDKN2A* and *TP53*, were preferentially deleted, reinforcing the role of LOH in driving tumorigenesis. Interestingly, we also observed that when CND occurs in the oncogene loci, they typically are selected for deletion of the reference allele rather than the mutated KRAS p.G12C/D or PIK3CA p.E545K allele.

Our results suggest that the interaction between CNVs and mutations can be both synergistic and context-dependent, as seen with genes like *PIK3CA* and *TP53*, where different levels of co-occurring alterations may implicate differing selection and expression impacts across various cancer types. We also show that in lung cancer, patients stratified by carrying these driver gene mutations only, CNV only, and co-occurring CNV/mutations of *EGFR*, *KRAS* or *TP53* had different prognosis. These complex interactions underscore the need to consider both mutations and CNVs in cancer diagnostics and treatment strategies to improve precision oncology.

Our study has several limitations. Somatic mutation and CNV calls from AACR GENIE were derived from different center’s panel sequencing and tumor-only data, and permutation results that were not validated in TCGA may not be as robust. Clinical data obtained from the GENIE BPC project may be biased and enabled only exploratory analyses that require validation using prospective or clinical trial cohorts. More, this data was only available for lung and colorectal cancers in a limited set of patients and was limited in statistical power. Also, the survival impact of treatments for *EGFR* (monoclonal antibodies and tyrosine kinase inhibitors) was inconclusive and partially limited by the fact that the GENIE BPC clinical data do not specify treatment sequencing or whether the treatment was completed. Additionally, while CNVs are well-characterized herein for their effects on mutation allele frequency and transcription, their role in non-coding regions and how they may cooperate with non-coding mutation drivers^38^ remains poorly understood. Lastly, emerging single-cell and spatial genomic datasets could help characterize these co-occurrences at a higher resolution^39^. Our study does not provide functional validation to track how a genetically engineered driver mutation may lead to selective amplification or deletion, however, experimental studies have demonstrated this for selected alleles, such as the amplification of genetically engineered KRAS G12D mutant alleles in mouse models^9–11^.

In conclusion, this study highlights the importance of somatic CNVs in modulating the effects of driver mutations and the role of co-occurring genetic alterations in cancer progression. Our findings emphasize the need for integrated genomic analyses in cancer research and the potential for developing therapeutic strategies that target these complex interactions. Further investigation into the mechanistic basis of mutation-CNV interactions will be crucial for advancing precision oncology.

## METHODS

### TCGA Genomics Data

Based on 10,956 unique TCGA cases, we included 8,546 pass-QC samples that had available CNV and mutation data.

Somatic driver mutations: Somatic mutations of 10,244 cases were obtained from the Multi-Center Mutation Calling in Multiple Cancers (MC3) dataset^24^. We only considered the nonsynonymous mutations in 299 cancer driver genes as defined by the PanCanAtlas driver project^40^, including missense, non-sense, frameshifting, in-frame shifting, or splice-site altering single-nucleotide changes or indels. Missense mutations predicted as having functional impact by any of the algorithms described in Bailey et al., as well as truncations, were considered as somatic driver mutations for analyses^41,42^.

Copy number variations: We obtained CNV deletions and amplifications of 9,125 TCGA cases as defined by the PanCanAtlas project^30^. Copy number values for each genomic segment were determined by applying ABSOLUTE to somatic DNA copy number data from 10,552 TCGA samples, and totals were determined by summing the number of detected segments within a given sample. The gene-level events indicate that the copy number gain/loss affects the specific genomic region that encodes the gene. CNV was assessed with Affymetrix SNP 6.0 arrays and gene-level CNV values were generated by GISTIC2^37^. GISTIC calls of −“2” and “2” which indicate a loss or gain of more than half of baseline ploidy were assigned as deep deletions or amplification, respectively.

Read count and VAF data for mutations: We processed DNA/RNA-seq BAM files from The Cancer Genome Atlas (TCGA), aligned to the GRCh38 reference genome, to analyze TCGA somatic mutation data provided in a mutation annotation format (MAF) file^43^. To harmonize the mutation coordinates with the RNA-seq data, we used the ‘vcf-liftover’ tool to remap the mutations to GRCh38, where all mutations were successfully lifted over. We then employed the ‘basecounts’ tool (https://github.com/PoisonAlien/basecounts) to calculate the base counts at each mutation site using the GRCh38 reference genome.

### AACR Project Genie (version 15.1)

Based on 171,145 unique cases provided by AACR Genie^25^ (v15.1), we combined mutation, CNV and clinical data from participating centers. It includes separate files of mutations, CNV, and clinical data, all keys by patient identifiers. For each cancer, a separate file of patient identifiers is created. For our analysis all these files were combined to create data sets by gene, cancer and other parameters.

### Identify significant co-occurring CNVs and mutations under selection

To determine if mutations and CNVs co-occur more often than expected by chance, we performed a permutation test to determine if the observed rate of mutation/CNV co-occurrence was greater than expected if there was no relationship between them. This framework is expanded upon the permutation analyses used for composite mutations^5,44^. We created matrices of CNA and CND observed mutation counts by sample (row) and gene (column). All such matrices containing a count of mutations by sample and gene, will be referred to as ‘mutation matrices ’in the method detailed below. For each cancer type in TCGA or the AACR Genie cohorts, the process is as follows:

#### Calculate the Observed Proportion

Create an observed mutation-only mutation matrix for that cancer type (matrix ‘MUT’) and an observed CNV mutation matrix (matrix ‘CNV’). These are then filtered only to contain genes and samples common to both matrices.

1. Find the observed proportion of mutations for each gene present in both MUT and CNV matrices and calculate gene mutation co-occurrence as a proportion. For example, if 2 out of 5 samples for Gene A have mutations present in both CNV and MUT, the Gene A gene mutation co-occurrence proportion is 2/5 = 0.4.
2. Remove from MUT and CNV matrices any genes with zero **observed** co-occurrence proportion. This means we have no data for this gene.
3. The result is a gene/mutation co-occurrence proportion for each gene in the MUT and CNV matrices.

Every CNV matrix was split into CNA and CND matrices, and every analysis was separately performed on CNA and CND matrices.

#### Calculate the Expected Proportion

The expected proportion of gene mutation co-occurrence was calculated by generating 100,000 random permutations of MUT and CNV matrices:

1. For each randomly generated MUT and CNV matrix, calculate gene mutation co-occurrence proportion. This will create an ALL_PERMUTATIONS table with 100K rows, for each gene common to MUT and CNV matrix.
2. For each gene in the ALL_PERMUTATIONS table, determine the ratio of permutations where the permuted proportion exceeds the observed proportion. This is the probability (after FDR correction) of mutation/CNV co-occurrence, assuming there is no relationship between them. This dataset is then saved to a file for each cancer type.

The GENIE permutation analysis was the same as the TCGA analysis, but the data was also partitioned by the participating center. Each center always provided data on a subset of genes and cancer. The permutation data for TCGA and each of the GENIE centers were combined, and multiple test statistics for single gene-cancer type pairs were combined using Edgington’s Method, to account for any zero probabilities generated by the permutation algorithm.

### Determining Significant Gene-cancer type pairs with Co-occurring CNV and Mutations

To evaluate the influence of copy number variations (CNVs) on mutation allele frequencies (VAFs) of driver mutations within each gene-cancer pairing, we implemented a multiple linear regression analysis. The model is formalized as follows:

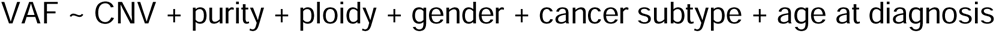

#### The Algorithm

1. From all the distinct gene cancer pairs select all those with enough mutations, with and without CNV, all with read depths of 5 or greater.
2. Split the dataset into:

a. pairs with mutations and CNV amplifications (CNA list)
b. pairs with mutations and CNV deletions (CND list)
3. For each list:

a. Construction and evaluation of a multiple regression model for each gene-cancer pair, noting the slope and significance of the CNV variable, which is invariably included as an independent variable across all models. CNV levels are quantified on a scale from -2 to +2, following the GISTIC 2.0 guidelines, where -2 indicates a deep deletion, 0 signifies mutation presence without CNV, +1 some amplification, and +2 denotes a high gain amplification.
b. Apply the Benjamini-Hochberg procedure to all p-values, to control the false discovery rate (FDR)

### Identifying Mutations Showing Allelic Imbalance at the DNA/RNA Levels

We developed the algorithm, Binoculars (Binomial test for co-occurring mutation and CNV pairs), was used for both RNA/DNA analyses. The significance of each specific mutation was determined by testing the read count number of wild-type reads and mutation-containing reads using exact binomial tests. The determination of the probability of success in the text was calculated using an algorithm where each mutation was matched against a background rate based on its most related data set:

For each gene-cancer pair, the data sets were searched in the following order, for the first match:

1. Find a match by (gene, cancer, and sample) in mutation-only data aggregated by gene, cancer, and sample
2. Find a match by (sample) in mutation-only data aggregated by sample only
3. Find a match by (gene, cancer) in mutation only data aggregated by gene and cancer
4. Find a match by (cancer) in mutation-only data aggregated by cancer only

If a match was found, this was the background rate parameter used for the binomial test’s probability of success. The Benjamini-Hochberg procedure was applied to all p-values, to control the false discovery rate (FDR).

### Transcriptional Effects of Co-occurring Mutations and CNVs

We focused on the likely driver mutations of the TCGA PanCan Atlas-identified 299 driver genes^40^ for this analysis using TCGA CNV, mutation, and gene expression data described above. The analyses were separated based on truncation vs. missense mutations. To identify co-occurring mutation and CNV that show synergistic effects on gene expression, we took a two-step approach:

1. We applied AeQTL^36^ to identify, for each cancer type, genes whose mutations are associated with expression upon adjusted for covariates (age, gender, ethnicity, cancer subtype [when available]) in multiple regression models where the dependent variable is expression and independent variable is the mutation status. The results were multiple-testing corrected using the Benjamini-Hochberg (BH) procedure for false discovery rate (FDR).
2. For the significant gene-cancer type pairs (FDR < 0.05), a linear regression model was fitted:

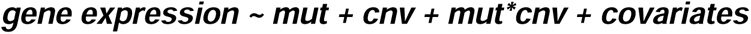

where mutation is the binary genotype (1 is mutated and 0 is wild type), gene-level CNV is a continuous variable (-2, -1, 0, 1, 2), mut*CNV is the interaction term, and the covariates are the same as described above. The regression coefficients of the interaction and its significance, after multiple testing corrections to FDR, were used to determine significant synergistic effects of co-occurring mutations and CNV on gene expression.

### eccDNA Analysis

Since the mutation data and eccDNA data^28^ used different assemblies (HG19 and HG39 respectively), we first performed a liftover of the eccDNA data to HG19. We obtained the HG19 gene positions from the UCSC Genome Browser (https://genome.ucsc.edu/index.html). We exported the known canonical data from the UCSC genes track. Using these gene positions, we performed a liftover on ECC gene data from HG38 to HG19. For each mutation’s gene, we found matching eccDNA fragments both long enough and which contain the entire gene (extrachromosomal, full-gene DNA). For each gene cancer pair, we counted the number of samples with mutations and CNV, the number of samples with genes found in eccDNA, and the number of samples with both.

Given the considerable variability in eccDNA length, ranging from hundreds to millions of base pairs, our analysis targeted identification of full gene eccDNA samples. These full gene eccDNA forms, prevalent in tumor cells, exhibit substantial size, enabling them to encapsulate fully functional gene copies.

### Survival Analysis

The Cox survival analysis was performed on mutation, CNV and clinical data for both lung and colorectal cancer. We performed analyses on key gene-cancer type pairs for both lung and colorectal cancers, and a single analysis for patients with EGFR mutations treated with combinations of tyrosine kinase inhibitors, monoclonal antibodies targeting EGFR receptors, and immunotherapy.

The covariates were age at diagnosis, stage, gender and mutation/CNV status (no mutations or CNVs, mutations only, CNVs only, both mutations and CNVs). The reference levels were “no mutations or CNVs” for CNV status, and “females” for gender. A patient was considered to have been treated with a drug if that drug was present in the drug data, regardless of treatment sequence, or how often the drug was administered during treatment cycles. There was insufficient detail in the data to analyze treatment sequencing effects, treatment outcomes or treatments abandoned due to side effects or complications.

All cox models were reviewed to ensure there were no proportional hazard assumption (PHA) violations (Schoenfield residuals) and ensure there were convergence. The single exception to this for the analysis on EGFR treated patients over the age of 60, where the monoclonal antibody (mab) covariate PHA was violated. Since this covariate’s impact was non-significant, and the violation was not severe (Schoenfield plot showed no significant relationship with survival time), it was retained in the model.

## Supporting information

Cover letter after 2nd response to reviewer

Cover letter after 1st response to reviewer

## DATA AND SOFTWARE AVAILABILITY

### Data Availability

These are the data files used in the analysis and detailed instructions on how to structure your file system is detailed in https://github.com/Huang-lab/CoMutCNV/DATA.md

| File | Source | Description |
| --- | --- | --- |
| aggregated_results.tsv | DNA/RNA-Seq readcounts extracted via <sup>43</sup> | DNA/RNA read count data of characterized mutations |
| all_thresholded.by_genes_whitelisted.tsv.gz | https://www.synapse.org/Synapse:syn22889333 | TCGA CNV Data |
| clinical_PANCAN_patient_with_followup.tsv | https://gdc.cancer.gov/about-data/publications/pancanatlas | TCGA PanCanAtlas patient data |
| hg19_canonical_gene_positions.csv | https://genome.ucsc.edu/index.html | Genome build data |
| mc3.v0.2.8.PUBLIC.maf.gz | https://gdc.cancer.gov/about-data/publications/pancanatlas | TCGA mutation data |
| subtype_assignments.csv | from [https://www.cell.com/cell/fulltext/S0092-8674(18)30237-X](https://www.cell.com/cell/fulltext/S0092-8674%2818%2930237-X) | TCGA PanCanAtlas patient cancer types |
| TCGA_mastercalls.abs_tables_JSedit.fixed.csv | https://gdc.cancer.gov/about-data/publications/pancanatlas | TCGA ABSOLUTE purity/ploidy file |
| driver_gene_by_cancer.txt | From TCGA PanCanAtlas Driver paper: [https://www.cell.com/cell/fulltext/S0092-8674(18)30237-X](https://www.cell.com/cell/fulltext/S0092-8674%2818%2930237-X) | driver gene classification by cancer type |
| All ATAC-seq files | http://genome.bioch.virginia.edu/TCGA-ATACSEQ-ECCDNA/ | TCGA EccDNA Data |
| AACR Genie | https://www.aacr.org/professionals/research/aacr-project-genie/aacr-project-genie-data/ | AACR Genie genomic data sets |
| AACR GENIE BPC | https://www.aacr.org/professionals/research/aacr-project-genie/bpc/ | AACR Genie clinical data sets |

### Code Availability

The source code is available at https://github.com/Huang-lab/CoMutCNV

## ACKNOWLEDGMENTS

The authors wish to acknowledge TCGA, AACR-Genie, and its participating patients and families that generously contributed the data. The authors thank all members of the Huang lab for constructive discussion. Large language models (LLM) may have been used in the initial drafts of coding and writing of this work. All final codes and texts have been extensively edited and verified by the authors. This work was supported in part through the computational and data resources and staff expertise provided by Scientific Computing and Data at the Icahn School of Medicine at Mount Sinai and supported by the Clinical and Translational Science Awards (CTSA) grant UL1TR004419 from the National Center for Advancing Translational Sciences. Research reported in this publication was also supported by the Office of Research Infrastructure of the National Institutes of Health under award numbers S10OD026880 and S10OD030463. The content is solely the responsibility of the authors and does not necessarily represent the official views of the National Institutes of Health. This work was supported by NIH NIGMS R35GM138113, NIGMS 2R35GM138113, ACS RSG-22-115-01-DMC, and Mount Sinai funds to KH.

## CONTRIBUTIONS

K.H. conceived the research, and K.H. and S.M. designed the analyses. S.M. developed the software. S.M., M.B., G.D., A.C., and Y.N-E conducted bioinformatics analyses based on H.C. and K.H. feedback. K.H. supervised the study. All authors read, edited, and approved the manuscript.

---

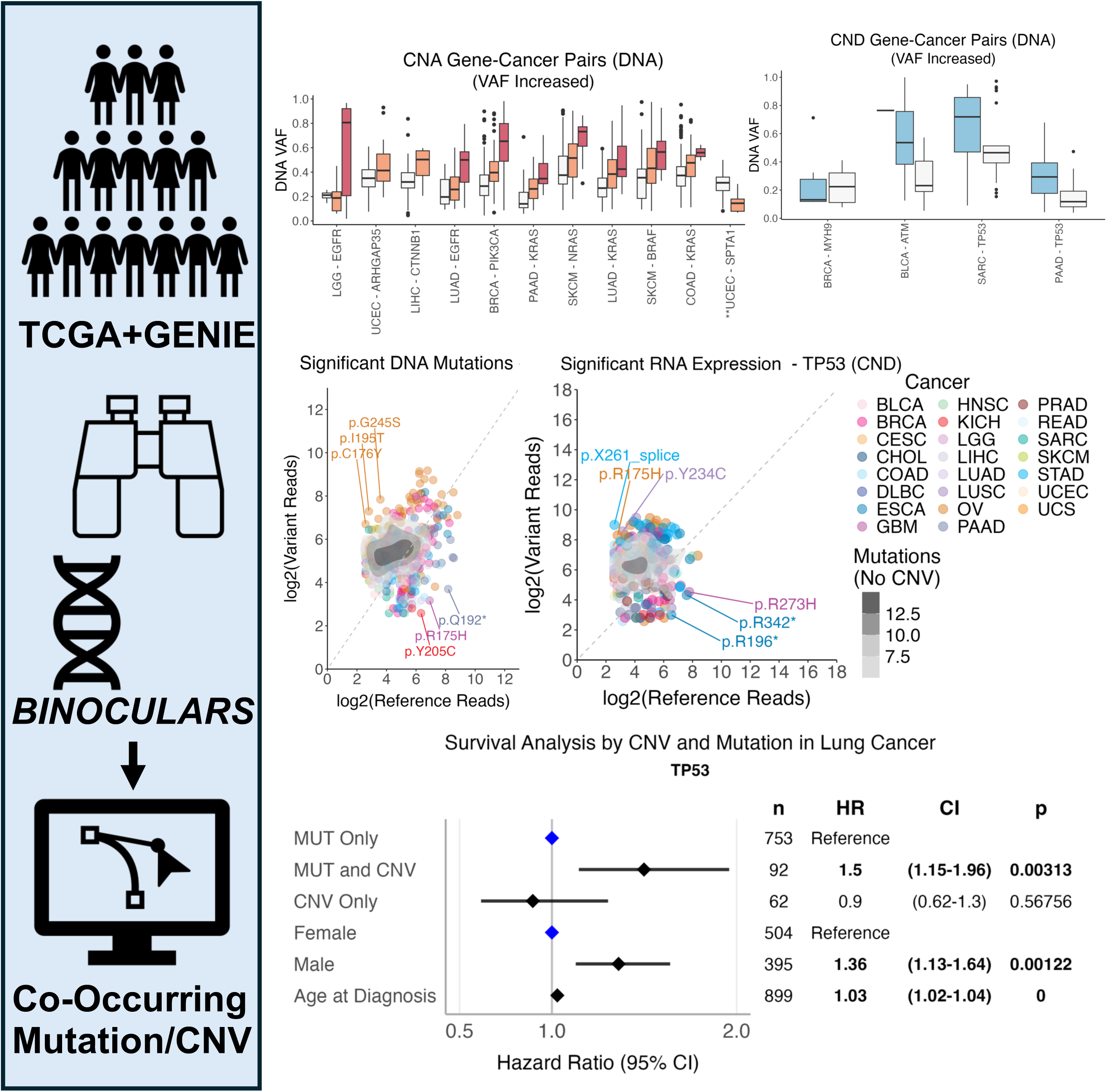

