## Supplementary figures and images for "Widespread Epistasis between Cancer Driver Mutations and Allele-Specific Copy Number Variations"

### Supplementary Figures

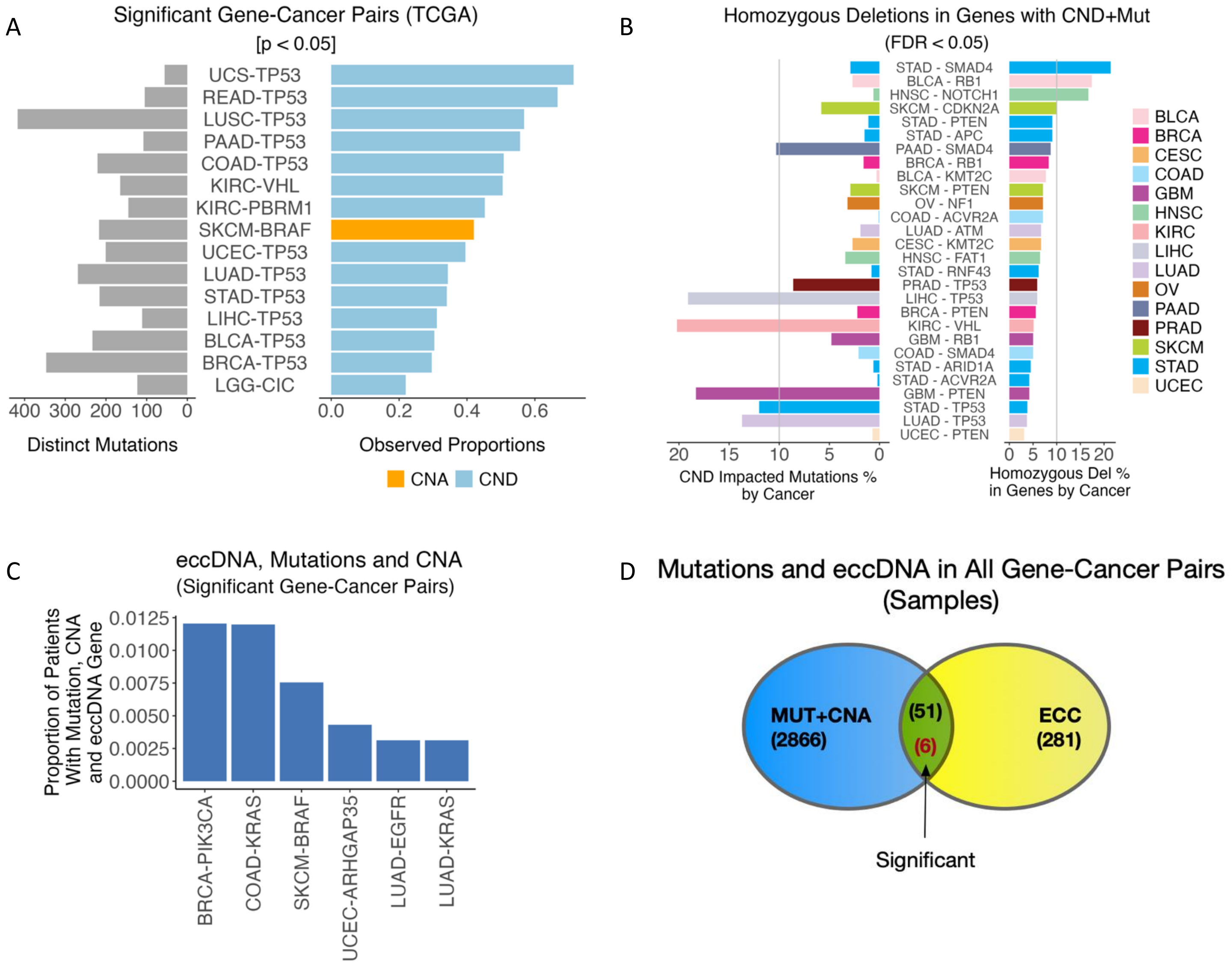

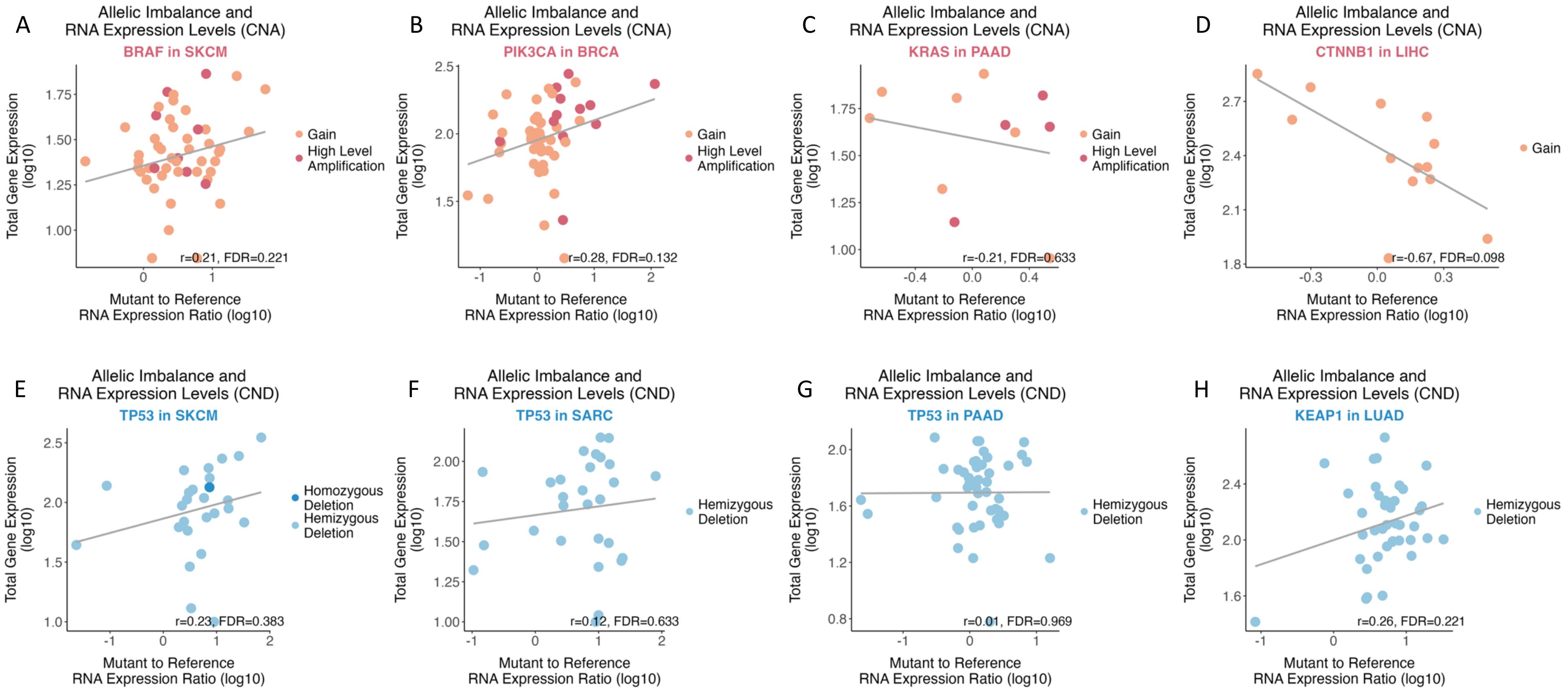

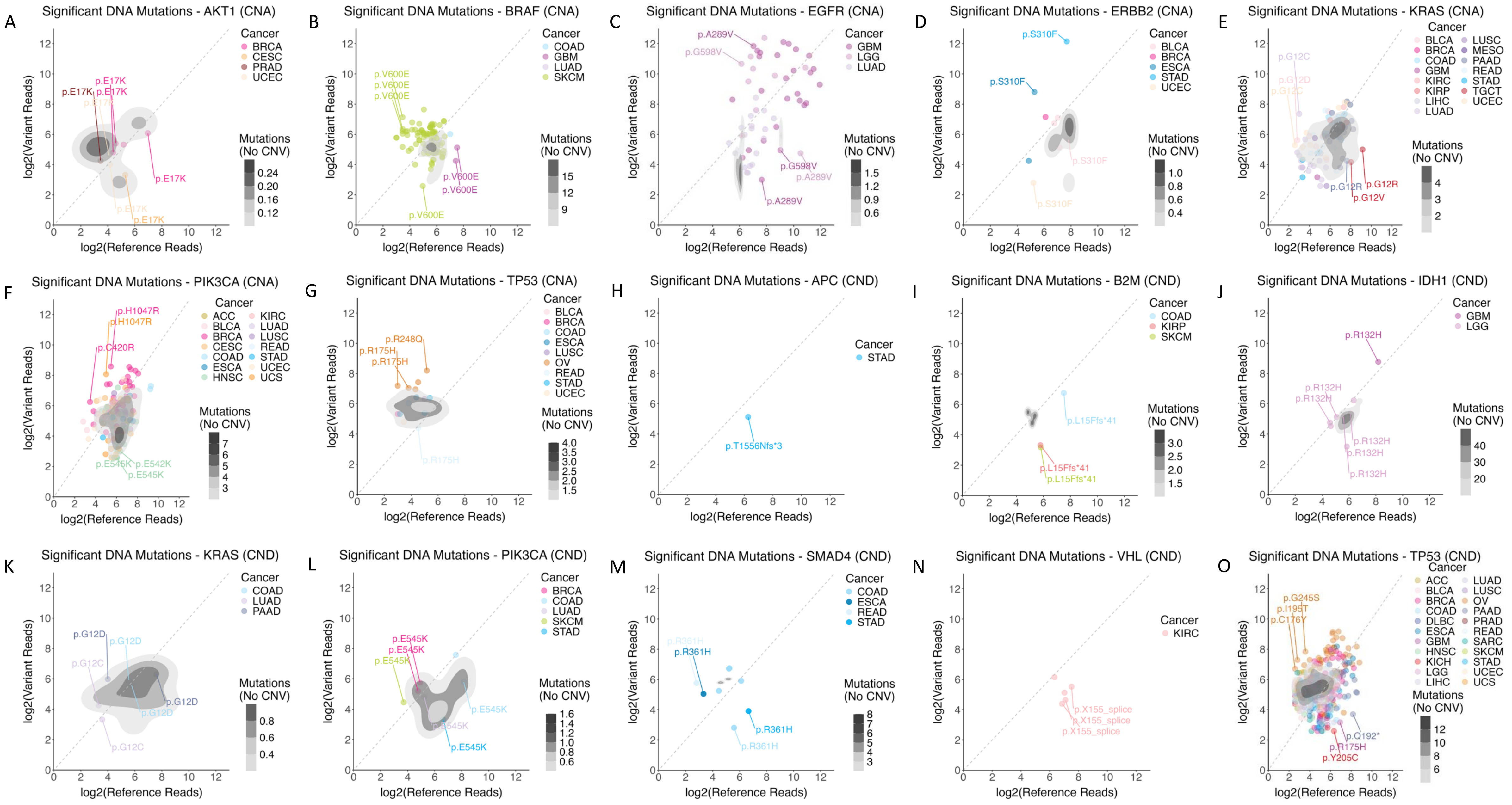

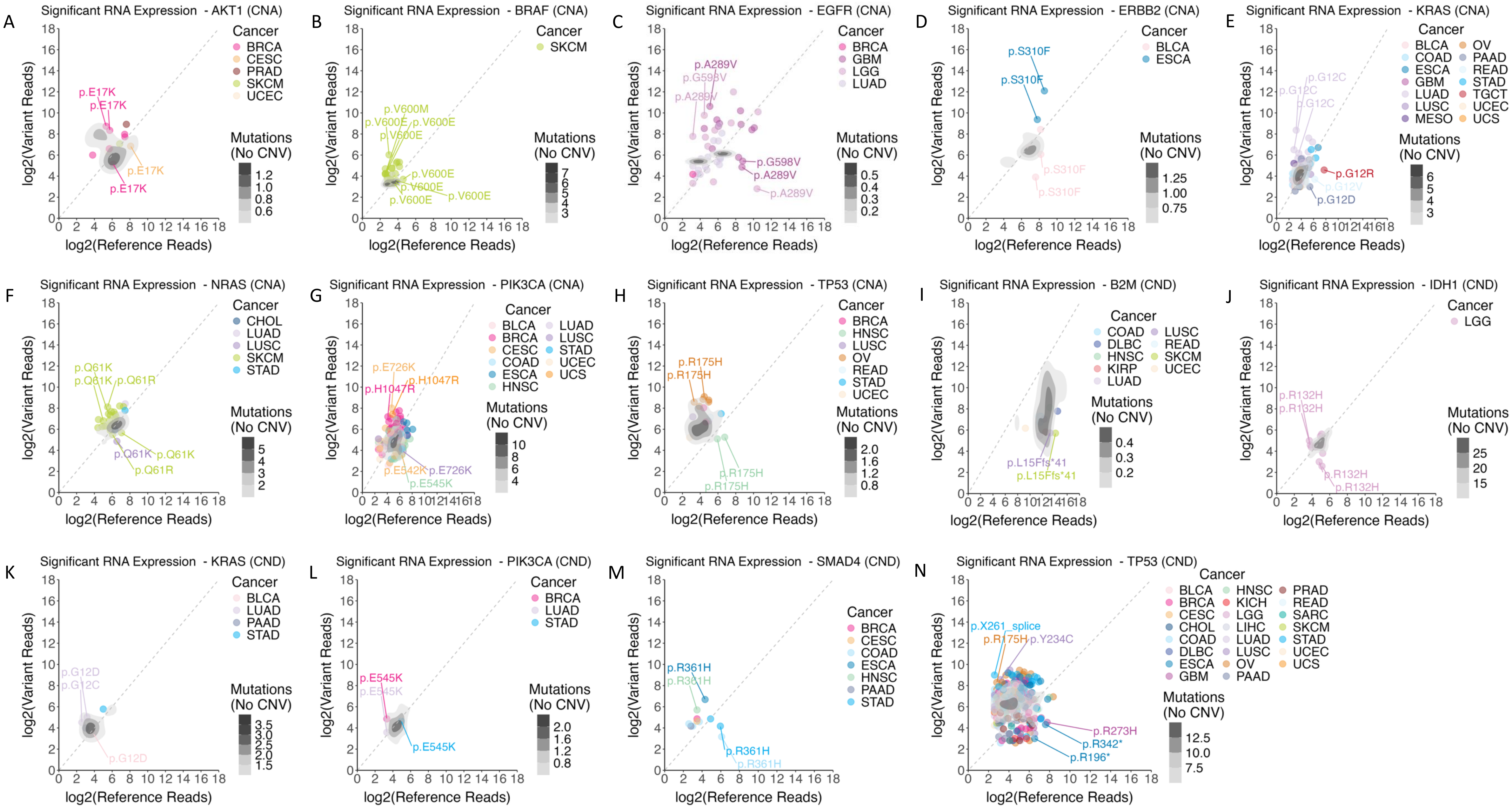

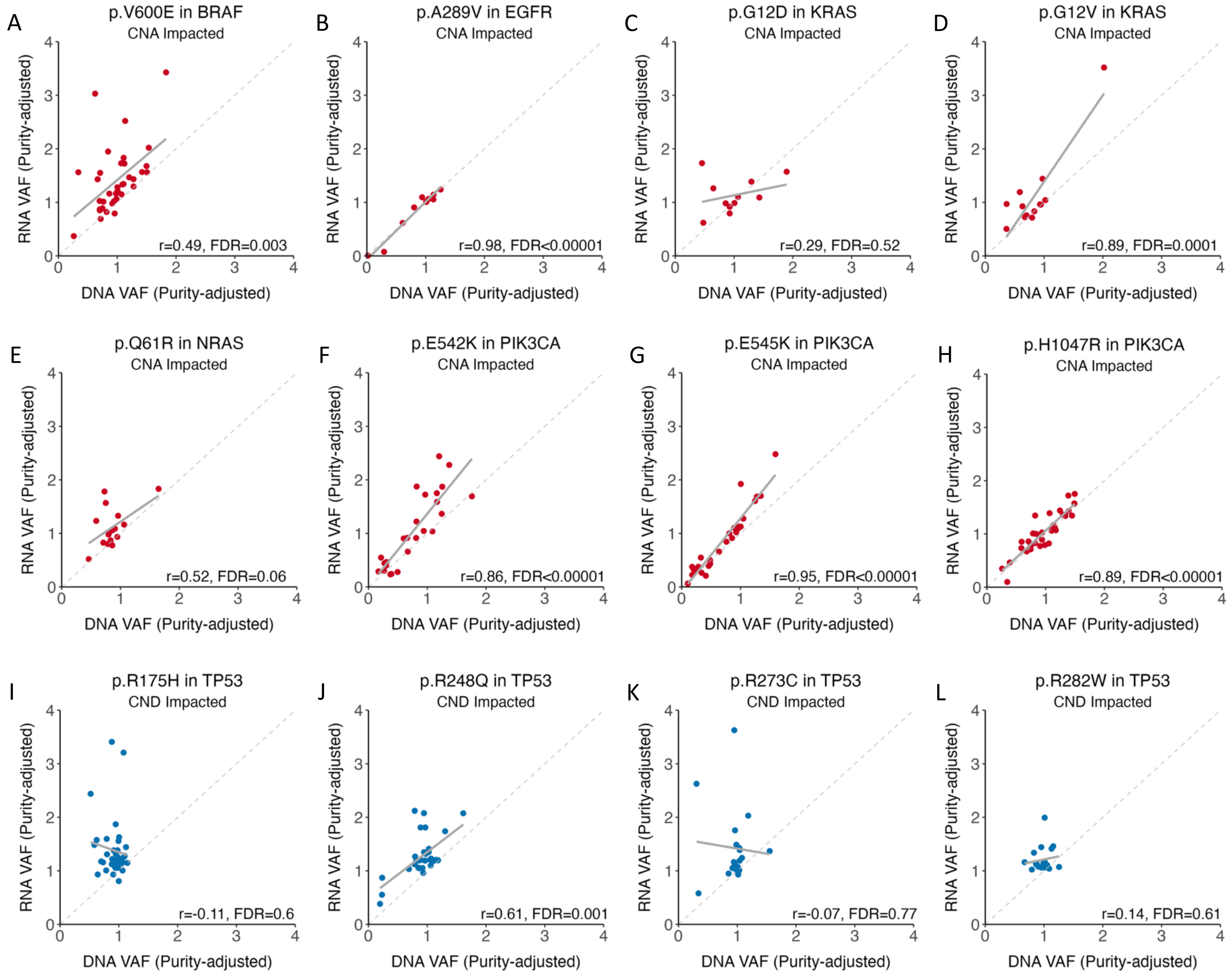

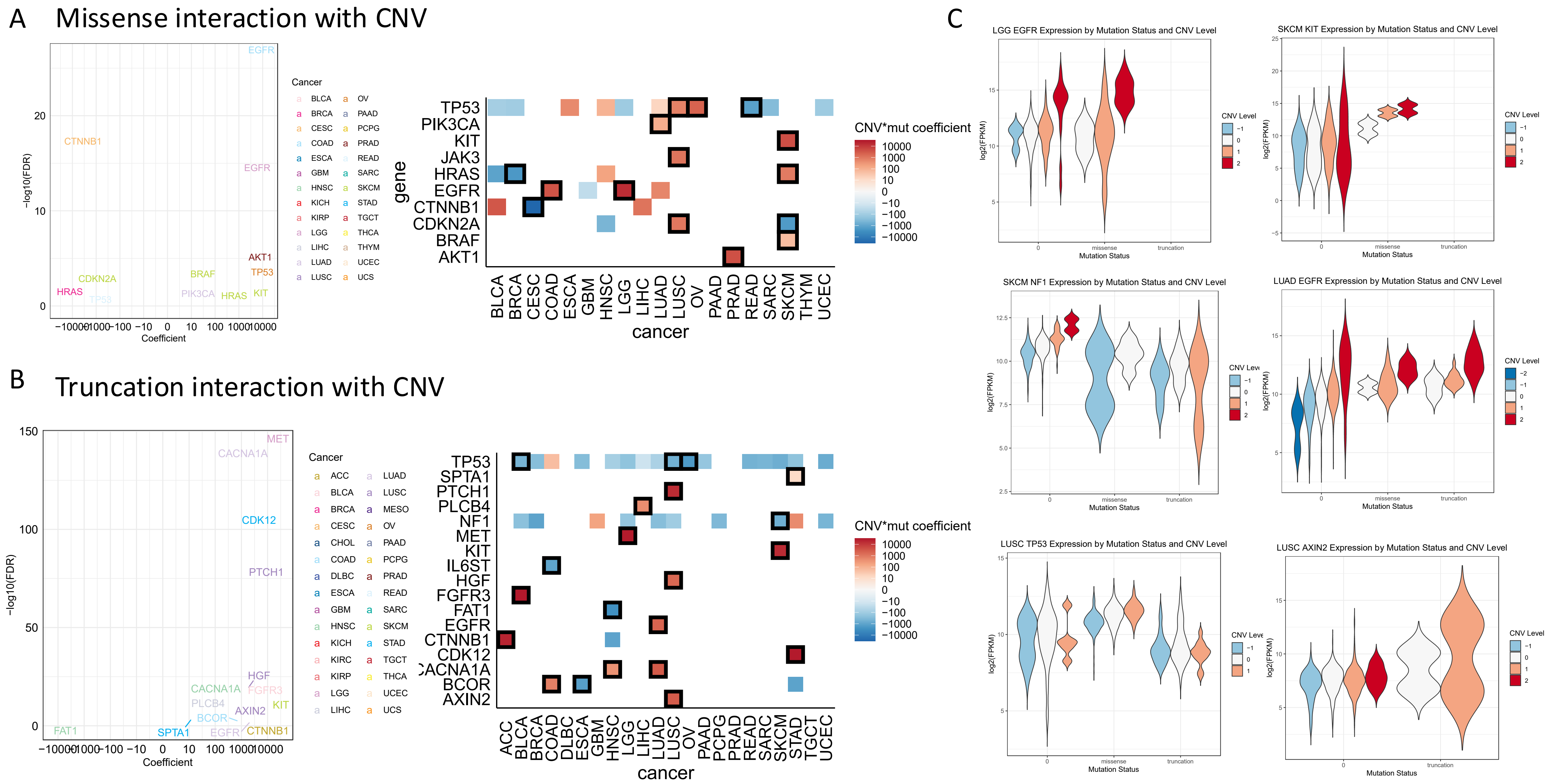

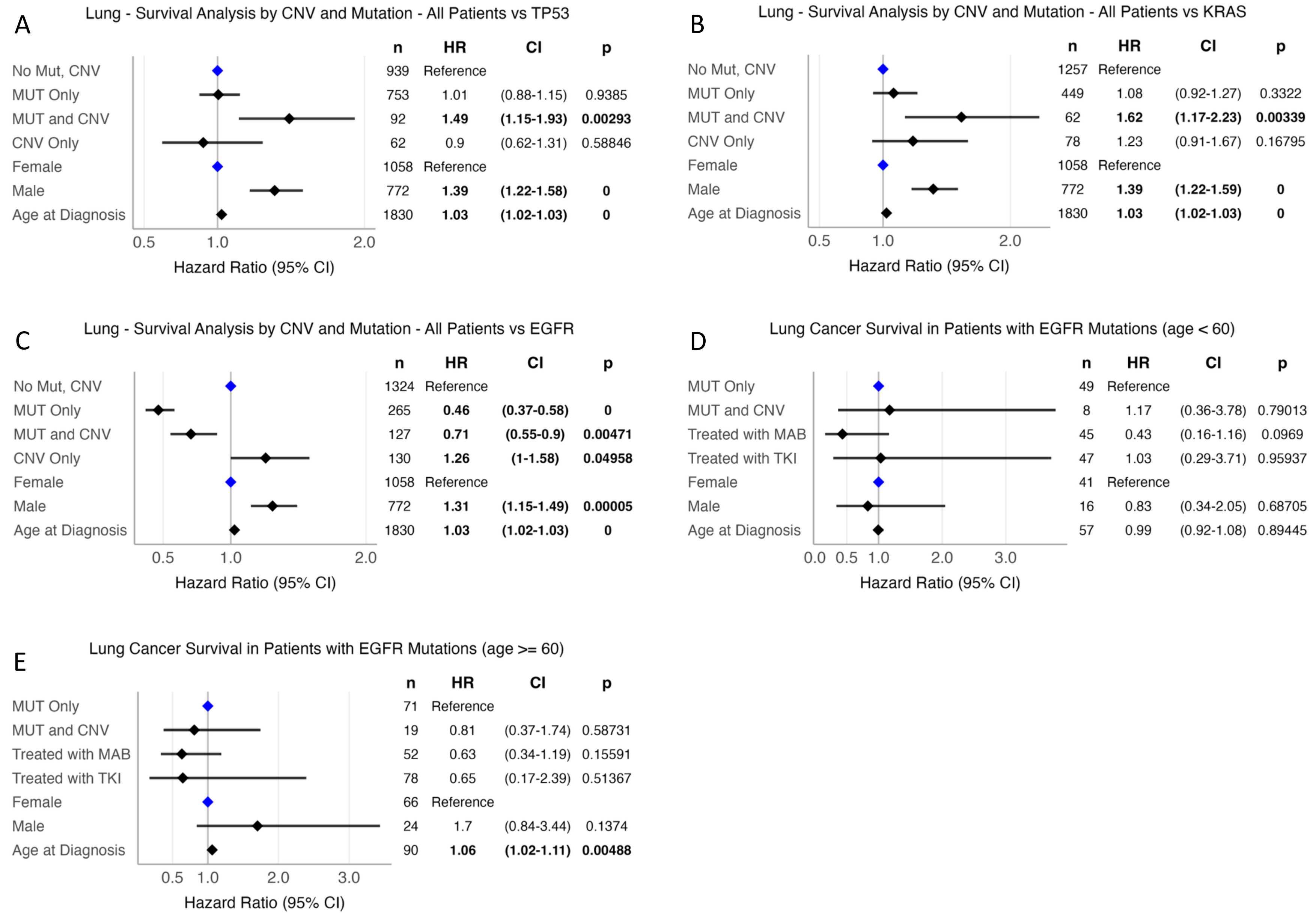

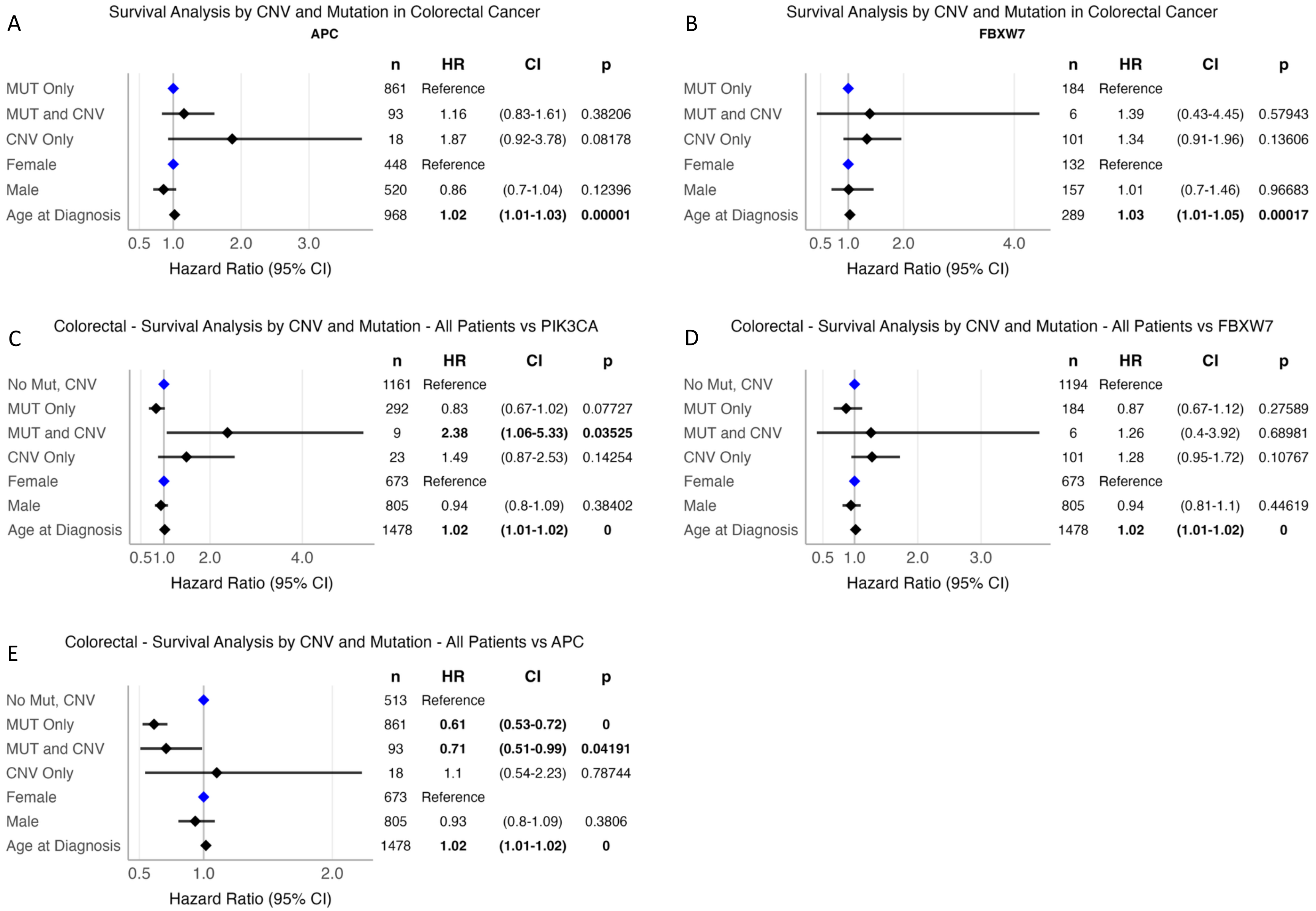
