## Supplementary material for "Widespread Epistasis between Cancer Driver Mutations and Allele-Specific Copy Number Variations": Cover letter after 2nd response to reviewer

Nov 30^th^ 2025

Luca Gasparoli, Ph.D.

Scientific Editor, Cell Reports

Dear Dr. Gasparoli,

Thank you again for handling our manuscript “Widespread Epistasis between Cancer Driver Mutations and Allele-Specific Copy Number Variations ” (CELL-REPORTS-D-25-04639R1). In light of Reviewer #2’s second-round report, we respectfully ask that you re-consider editorial decision at Cell Reports.

In this second-round review, most of the remaining concerns appear to reflect points that are already addressed and may have been misunderstood. We cite the reasons below and detailed point-by-point response are in the attached rebuttal. In brief:

- **We do not claim CNV–mutation epistasis is uniquely required**, but instead test whether specific driver mutations–CNV pairs are under selection, which in our knowledge is the first study to be done at this scale.
- **The Supek work was cited in the original submission** and it, like several other recent works, provided DNA interaction of mutation+CNVs at gene-level resolution. Our contribution uniquely provides mutation- and allele-resolved DNA and RNA analyses plus expression and clinical outcome associations.
- For TP53 p.R275, our results **reproduced the expected LOH patterns while adding rarer, mutant-allele–biased amplifications**, in line with known biology that the reviewer expected to see while also illuminating lesser-known biology.
- For key hotspots (including TP53), DNA and RNA allelic imbalance are **concordant** **in direction** but we chose to highlight the imperfect correlation to suggest the added insights provided by RNA level analyses.

Because these clarifications directly address the reviewer’s conceptual objections without changing the core, internally consistent results with fully reproducible data and code, we believe the manuscript now meets the bar for publication in *Cell Reports*. We would be very grateful if you could consider acceptance on this basis.

Sincerely and on behalf of the team,

Kuan-lin Huang, Ph.D.

Associate Professor of Genetics and Genomic Sciences & Artificial Intelligence and Human Health

Icahn School of Medicine at Mount Sinai

New York, NY 10029
