## Supplementary material for "Widespread Epistasis between Cancer Driver Mutations and Allele-Specific Copy Number Variations": Cover letter after 1st response to reviewer

Sep 30^th^ 2025

Luca Gasparoli, Ph.D.

Scientific Editor, Cell Reports

Dear Dr. Gasparoli,

We are pleased to submit our revised manuscript entitled *“Widespread Epistasis between Cancer Driver Mutations and Allele-Specific Copy Number Variations”* to *Cell Reports*, along with a detailed point-by-point response to the reviewers’ comments.

Following your suggestions and the reviewers’ feedback, we have conducted several additional analyses and made substantial revisions that significantly strengthen the manuscript. Here are the key improvements in this revised version:

1. We corrected factual discrepancies between text and figures and fully updated the Results, figure callouts, and conclusions to match the revised analyses, and updated all analyses to use FDR < 0.05. We also provide Figure 1A to clarify use of data for each analysis/figure.
2. We conducted new analyses to directly link allelic imbalance with total gene expression, stratified *TP53* by mutation type, added purity-adjusted VAFs, comparing homozygous vs. co-occurring mutation/CNV two-hits, and correlating DNA vs. RNA VAFs.
3. We expanded discussion and figures to address key reviewer points, including homozygous deletions vs. mutation-CNV “two-hit” events, GENIE data limitations, discordant DNA/RNA allelic imbalance, mutation vs. CNV timings that may result in allelic imbalance, and added reference while clarifying our innovations of identifying CNV-selected mutations at a mutation (rather than gene) resolution.

Additionally, we carefully edited the manuscript for accuracy, improved figure clarity, and expanded the discussion. We believe the revised manuscript adequately addresses all reviewer concerns and provides a clearer and more comprehensive analysis. We thank you and the reviewers for the thoughtful feedback and hope that you will find the revised manuscript suitable for publication in *Cell Reports.*

Sincerely and on behalf of the team,

Kuan-lin Huang, Ph.D.

Associate Professor of Genetics and Genomic Sciences & Artificial Intelligence and Human Health

Icahn School of Medicine at Mount Sinai

New York, NY 10029
